# Horizontal transfer of microbial toxin genes to gall midge genomes

**DOI:** 10.1101/2021.02.03.429655

**Authors:** Kirsten I. Verster, Rebecca L. Tarnopol, Saron M. Akalu, Noah K. Whiteman

**Author notes:** **Corresponding author**: Dr. Noah K. Whiteman, Department of Integrative Biology, University of California, Berkeley, Berkeley, CA 94720. equal contribution (Kirsten I. Verster, Rebecca L. Tarnopol).

## Abstract

A growing body of evidence points to a role for horizontal gene transfer (HGT) in the evolution of animal novelties. Previously, we discovered the horizontal transfer of the gene encoding the eukaryotic genotoxin cytolethal distending toxin B (CdtB) from the *Acyrthosiphon pisum* Secondary Endosymbiont (APSE) bacteriophage to drosophilid and aphid genomes. Here, we report that *cdtB* is also found in the nuclear genome of the gall-forming ‘swede midge’ *Contarinia nasturtii* (Diptera: Cecidomyiidae). We subsequently searched genome sequences of all available cecidomyiid species for evidence of microbe-to-insect HGT events. We found evidence of pervasive transfer of APSE-like toxin genes to cecidomyiid nuclear genomes. Many of the toxins encoded by these horizontally transferred genes target eukaryotic cells, rather than prokaryotes. In insects, catalytic residues important for toxin function are conserved. Phylogenetic analyses of HGT candidates indicated APSE phages were often not the ancestral donor of the toxin gene to cecidomyiid genomes, suggesting a broader pool of microbial donor lineages. We used a phylogenetic signal statistic to test a transfer-by-proximity hypothesis for HGT, which showed, that prokaryotic-to-insect HGT was more likely to occur between taxa in common environments. Our study highlights the horizontal transfer of genes encoding a new functional class of proteins in insects, toxins that target eukaryotic cells, which is potentially important in mediating interactions with eukaryotic pathogens and parasites.

**Significance Statement:** The diversity of genes encoded by phages infecting bacterial symbionts of eukaryotes represents an enormous, relatively unexplored pool of new eukaryotic genes through horizontal gene transfer (HGT). In this study, we discovered pervasive HGT of toxin genes encoded by *Acyrthosiphon pisum* secondary endosymbiont (APSE) bacteriophages and other microbes to the nuclear genomes of gall midges (Diptera: Cecidomyiidae). We found five toxin genes were transferred horizontally from phage, bacteria, or fungi into genomes of several cecidomyiid species. These genes were *aip56, cdtB, lysozyme, rhs*, and *sltxB*. Most of the toxins encoded by these genes antagonize eukaryotic cells, and we posit that they may play a protective role in the insect immune system.

## Main Text

There is growing evidence that horizontal gene transfer (HGT) from microbes to animals has played an important role in animal evolution (Husnik & McCutcheon 2018). Toxin-encoding genes have been horizontally transferred into arthropod genomes and even integrated into their immune systems (Di Lelio et al. 2019; Hayes et al. 2020; Li et al. 2021). However, many of these events involve transfer of genes encoding toxins that target prokaryotes, and few characterized HGTs in animals are of genes encoding toxins that target eukaryotes.

We previously discovered HGT of a eukaryote-targeting toxin gene, *cytolethal distending toxin B* (*cdtB*), into the nuclear genomes of four insect lineages within two orders, Diptera and Hemiptera (Verster et al. 2019). The closest relatives of these insect *cdtB* copies were copies isolated from *Acyrthosiphon pisum* secondary endosymbiont (APSE) bacteriophage (Verster et al. 2019), which infect the secondary bacterial endosymbiont *Hamiltonella defensa* (Degnan & Moran 2008; Oliver et al. 2009, 2010) of hemipterans and other cosmopolitan symbiotic bacteria such as *Arsenophonus* spp. (Duron 2014). APSE phages encode diverse toxins within a highly variable “toxin cassette” region of their genomes (Rouïl et al. 2020). We found another APSE toxin, *apoptosis inducing protein 56* (*aip56)*, fused to an additional full-length *cdtB* copy in nuclear genome sequences of all *Drosophila ananassae* subgroup species examined (Verster et al. 2019). The synteny of *aip56* and *cdtB* genes is the same in APSE genomes, which further supports HGT of toxin genes from APSE phages to insect nuclear genomes.

Subsequently, we serendipitously discovered a full-length *cdtB* sequence in the nuclear genome sequence of the gall midge *Contarinia nasturtii* (Diptera: Cecidomyiidae) (**Table S1**). The Cecidomyiidae (Diptera: Nematocera) contains over 6,600 fly species with diverse life histories, behaviors and host use patterns (Yukawa & Rohfritsch 2005; Dorchin et al. 2019; O’Connor et al. 2019). Many cecidomyiids create destructive galls on crops (Hall et al. 2012). Previously, an APSE-3-like rearrangement hotspot (RHS) toxin gene was found in the genome of the wheat pest *Mayetiola destructor* (Zhao et al. 2015). To further investigate the extent of HGT from APSE to cecidomyiid genomes, we conducted tblastn searches using proteins encoded by APSE genomes as queries against all publicly available cecidomyiid whole genome sequences: *C. nasturtii, M. destructor, Sitodiplosis mosellana*, and *Catotricha subobsoleta* (**Table S2**). Each of these species had genomic reads and assembled contigs, and all but the latter had transcriptomic data available. We generated a short list of HGT candidates by excluding top matches to *bona fide* insect genes, hits <50 AA long, and hits on short scaffolds (for more details see **Supplementary Methods**).

We considered there to be strong evidence of HGT if the candidate gene of interest (GOI) met at least 2/5 of the following criteria:

1. Non-anomalous read depth via BWA analysis
2. The GOI is on scaffolds with other *bona fide* eukaryotic genes
3. The GOI is syntenic in two or more species
4. The GOI is transcribed in dT-enriched transcriptomes
5. The GOI is predicted to have introns

Many of these genes show some signatures of eukaryotic domestication (**Supplementary Text**). For *C. nasturtii*, we validated HGTs with PCR and bi-directional Sanger sequencing (see **Supplementary Methods** and **Table S4**) of genomic DNA from larvae and adults of this midge species. A summary of our QC methods for each species is shown in **Supplementary File 1** and our list of HGT candidates is shown in **Table 1**.

**Table 1.** Final list of HGT candidate genes from sequenced cecidomyiid genomes.

| Species | Protein Name | Scaffold(s) | GenBank ID or scaffold coordinates |
| --- | --- | --- | --- |
| *C. nasturtii* | Aip56 | NW_022197544.1<br>NW_022200251.1 | LOC116352557<br>LOC116349575 |
|  | CdtB | NW_022197544.1<br>NW_022203704.1 | LOC116352617<br>LOC116351853 |
|  | Lysozyme | NW_022201606.1 | LOC116350899<br>LOC116350797<br>NW_022201606.1: 2605394-2605828 |
|  | SltxB | NW_022197768.1 | LOC116338454<br>LOC116338453 |
| *S. mosellana* | Aip56 | VUAH01001532.1 | VUAH01001532.1: 2829-2410 |
|  | Hypothetical Protein | VUAH01004499.1<br>VUAH01000029.1 | VUAH01004499.1: 1380-1123<br>VUAH01000029.1: 4532498-4532776<br>4520662-4520964 |
|  | RHS | VUAH01000029.1 | VUAH01000029.1:<br>8-4950 |
|  | SltxB | VUAH01000060.1<br>VUAH01000016.1<br>VUAH01006166.1 | VUAH01000060.1: 423458-423673<br>425034-425252<br>VUAH01000016.1: 900502-900260<br>896597-896355<br>901304-901146<br>899522-899256<br>VUAH01006166.1: 4792842-4792618 |
| *M. destructor* | Aip56 | GL501523 | GL501523: 1201296-1200694 |
|  | Lysozyme | AEGA01007297.1<br>AEGA01024319.1<br>GL501497 | Mdes015794<br>Mdes01448<br>GL501497: 230410-230841 |
|  | RHS protein | GL501425 | Mdes011034 |

The short list of HGT candidates almost exclusively includes toxin genes. They are *aip56, cdtB, lysozyme, rhs toxin*, and *Shiga-like toxin B* (*sltxB*). Additionally, we found multiple copies of an APSE-4 hypothetical protein in *S. mosellana*. This gene is found in the “toxin cassette” of APSE genomes (Rouïl et al. 2020). We excluded it from further analyses as it is poorly characterized form a functional perspective. To discern the timing and provenance of these HGTs, we incorporated phylogenetic information and, where applicable, synteny information (see **Figure 1, Figure S1**, and **Table S3**). We then used structural analysis with Phyre2 (Kelley et al. 2015) and MAFFT (Katoh et al. 2019) to determine if they have retained their function following transfer into insect genomes. Below we summarize our findings for each of the HGT candidates.

**Fig 1.**
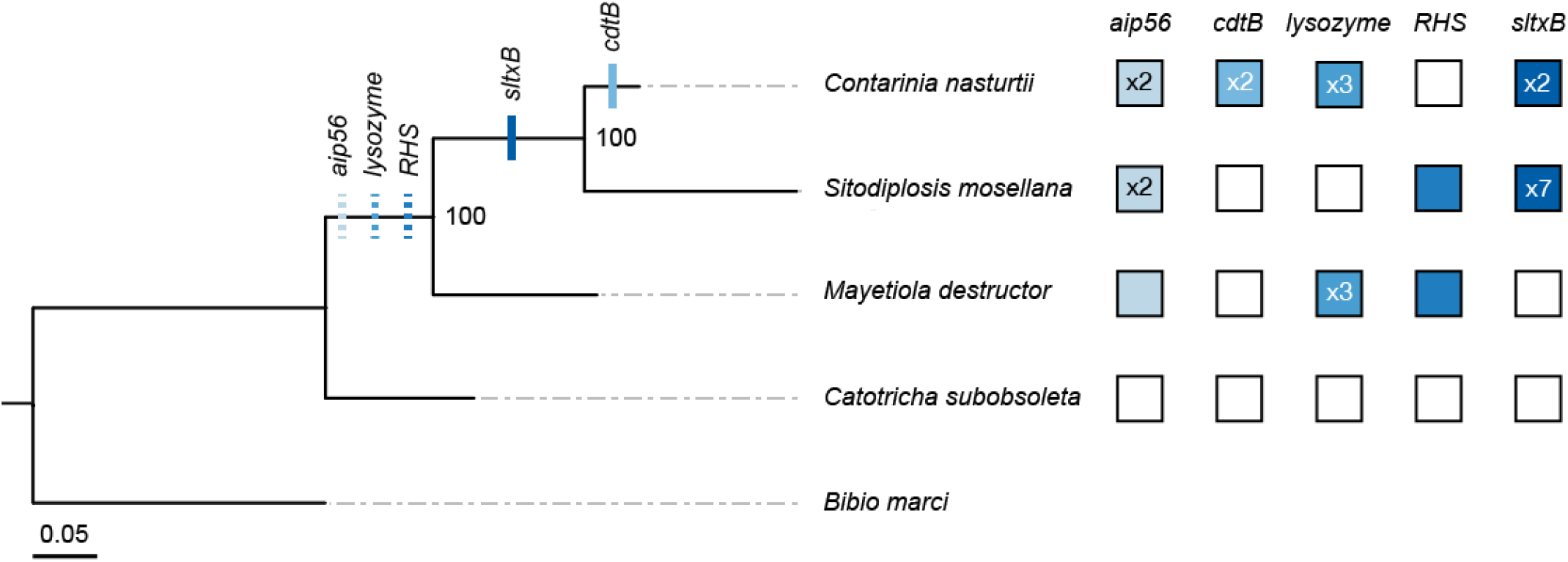
Maximum likelihood Cecidomyiidae species phylogeny (log likelihood = −10311.662040) shows the approximate timing of each HGT event. The tree was built in RAxML using concatenated sequences of the *co1* (542 nt), *CAD* (1439 nt), *ef1a* (725 nt), and *28S* (429 nt) genes (see **Table S7** for accession IDs for genes included in the species phylogeny). *Bibio marci* (Diptera: Bibionidae) is included as an outgroup. Filled boxes indicate presence of the toxin, and the numbers indicate copy number if >1. Bootstrap values are reported out of *n* = 1000 bootstraps, and scale bar is substitutions per site. Tick marks on the phylogeny indicate approximate timing of the HGT event based on a parsimony approach incorporating presence/absence of the HGT candidate, individual gene phylogenies, and synteny data. Dashed ticks indicate HGT events for which synteny data were inconclusive.

### Aip56

Aip56 is a secreted toxin of *Photobacterium damselae* subsp. *piscicida*, a fish pathogen that induces apoptosis of blood cells (do Vale et al. 2017). We previously found the Aip56 B domain encoded in a fusion gene comprised of a full-length *cdtB* copy and a partial *aip56* copy. The Aip56 B domain facilitates internalization to target cells (Pereira et al. 2014), and was horizontally transferred to the *Drosophila ananassae* species complex from an APSE-like phage (Verster et al. 2019).

Insect Aip56 protein sequences form a paraphyletic clade consisting largely of insects or insect symbionts (**Figure 2, Figure S1**). Cecidomyiidae Aip56 is closely related to sequences that include Lepidoptera-associated viruses and several other insect taxa (**Figure 2**), indicating *aip56* has been transferred multiple times in insects from the APSE phage lineage. As in previous studies (Silva et al. 2013; Verster et al. 2019), we did not find conservation of the zinc-binding motif HEXXH in insect or insect-associated sequences, so catalytic activity is likely absent in insect Aip56 (**Figure 3**). Short domains appear to be conserved in the Aip56 B domain (Verster et. al. 2019), which is necessary for cellular uptake of the toxin (Silva et al. 2013; Pereira et al. 2014). However, given the dearth of information about Aip56, it is difficult to assess their biological significance. Still, domains in insect Aip56 show homology to immunity proteins and lectin-binding motifs (**Table S5**), suggesting an immune function.

**Fig 2.**
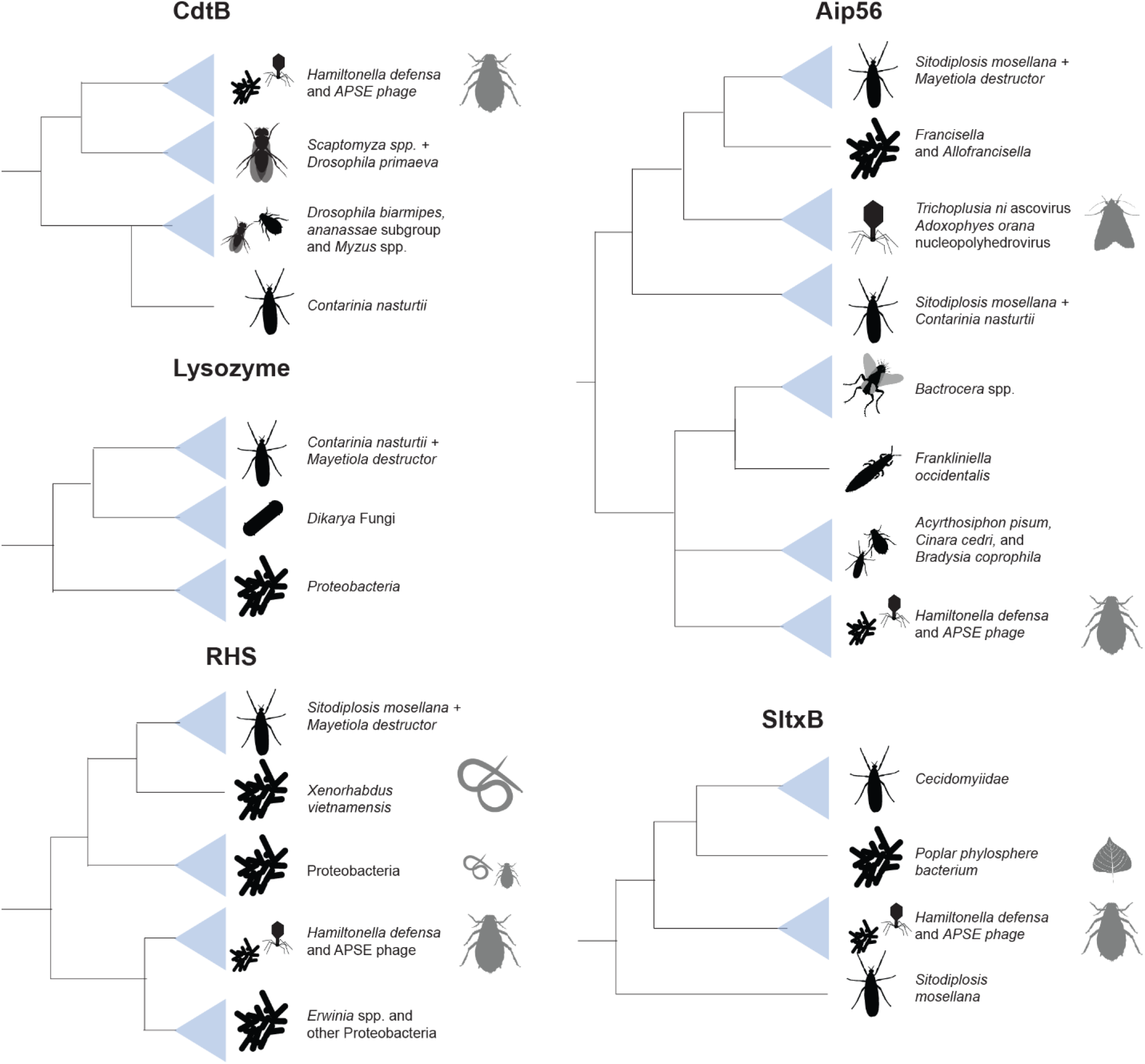
Simplified phylogenies of horizontally transferred genes show they are derived from several sources, including endosymbiont bacteriophage, Proteobacteria, and fungi. Indicated co-associated species suggest opportunities facilitating HGT events. Black organisms are species, while grey organisms are co-associated species. For full phylogenies, see **Figure S1**.

**Fig 3.**
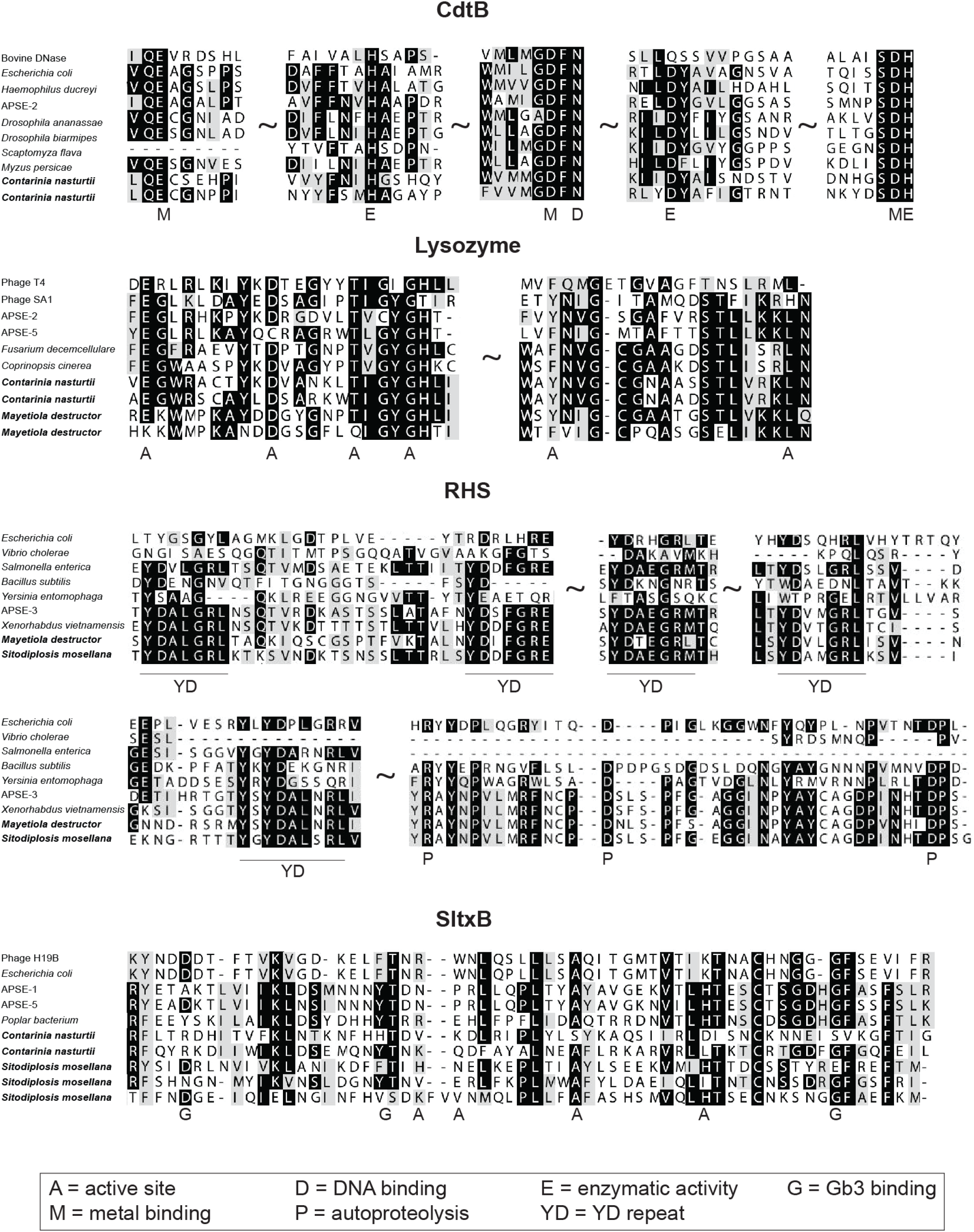
Protein alignments of toxin proteins transferred to midges reveal that many critical residues are conserved (see **Supplementary Methods** for accession IDs of representative sequences).

### CdtB

CdtB is a DNase I encoded within the genomes of diverse Actinobacteria and Proteobacteria (Jinadasa et al. 2011). CdtB complexes with CDT subunits A and C to enter eukaryotic cells, after which CdtB nicks the DNA, triggering mitotic arrest and apoptosis (Jinadasa et al. 2011). In aphids, CdtB plays a role in resistance of aphids to parasitoid wasps (Oliver et al. 2009), and we suspect it retains this function in drosophilids (Verster et al. 2019).

Since we only found *cdtB* in the genome of *C. nasturtii* among the gall midge genome sequences we searched, we infer that it was introduced into the genome after the split with *S. mosellana* ancestors ca. 70 mya (Dorchin et al. 2019). The CdtB phylogeny shows *C. nasturtii* CdtB is monophyletic with other insects, endosymbiotic bacteria, and phage (**Figure 2**, see also **Figure S1**), consistent with our previous study (Verster et al. 2019). We hypothesize that CdtB was donated from the genome of an endosymbiotic bacterium or the APSE phage lineage to a *C. nasturtii* ancestor. Consistent with previous analyses (Verster et al. 2019), amino acid residues important for CdtB metal binding, DNA binding, and enzyme activity (Jinadasa et al. 2011) were conserved in *C. nasturtii* (**Figure 3, Figure S5)**, indicating the ancestral function is maintained.

### Lysozymes

Lysozymes hydrolyze glycosidic bonds in peptidoglycan, a component of bacterial cell walls. Lysozymes play diverse roles including in immune defense, bacterial digestion, bacterial cell wall synthesis, and release of mature phages from infected bacterial cells (Van Herreweghe & Michiels 2012).

Interestingly, our phylogeny shows that Cecidomyiidae lysozyme genes were transferred from fungi, rather than from the APSE phage lineage. The cecidomyiid lysozyme sequences (*M. destructor* + *C. nasturti*i) are nested in a highly supported monophyletic clade sister to the fungal phyla Ascomycota and Basidiomycota, distant from APSE lysozyme sequences (**Figure 2, Figure S1**). This fungal lysozyme clade is sister to a large clade of lysozymes from Proteobacteria, consistent with the fact that GH25 lysozymes were transferred across the tree of life from Proteobacterial donors (Metcalf et al. 2014). The cecidomyiid and fungal lysozyme sequences share high structural similarity with phage lysozyme GH24 (**Table S5**). Previous studies show fungal GH24 lysozyme may play a role in nematode defense (Plaza et al. 2015; Kombrink et al. 2019). Many residues vital for binding and catalysis (Shoichet et al. 1995) are the same between insect, fungal, and phage lysozyme sequences, indicating a conserved proximal function in insects (**Figure 3**).

### RHS toxins

Rearrangement hotspot (RHS) toxins, or YD-repeat toxins, are found widely among bacteria and archaea (Jamet & Nassif 2015). RHS toxins are large and highly polymorphic, consisting of several tyrosine/aspartate (YD) repeats that are involved in trafficking and delivery of the toxin and a variable C-terminal domain that catalyzes the enzyme’s toxic activity (Zhang et al. 2012).

The cecidomyiid RHS proteins form a single clade sister to *Xenorhabdus vietnamensis*, a symbiotic bacteria of the entomopathogenic nematode *Steinernema sangi* (Lalramnghaki et al. 2017) (**Figure 3**). This clade, in turn, is sister to a group that includes *Xenorhabdus* and *Photorhabdus* species, which are associated with nematode parasitism of insects (Boemare 2002; Busby et al. 2013). The most inclusive clade includes APSE phage and associated endosymbionts. The APSE-3 RHS toxin has been implicated in parasitoid defense, as pea aphids harboring *H. defensa* infected with APSE-3 had near-complete protection against parasitism by the parasitoid braconid wasp *Aphidius ervi* (Martinez et al. 2018). The cecidomyiid RHS sequences retain residues important for toxin function. Insect RHS sequences maintain the YDXXGR core repeat motif shared among bacterial RHS toxins (Wang et al. 1998). Additionally, three residues involved in C-terminal autoproteolysis, R650, D663, and D686 (Busby et al. 2013), are conserved in insect RHS toxin copies (**Figure 3**). Cecidomyiid RHS toxins show structural similarity to the insecticidal *P. luminescens* Tc toxin complex (**Figure S8**), consistent with a toxic functional role.

### SltxB

Shiga-like toxins (Sltxs) are ribosome-inactivating toxins (Chan & Ng 2016). Sltxs are AB_5_ toxins, where the B pentamer binds to globotriaosylceramide (Gb3) binding sites to retrograde traffic the active A subunit into the cell (Malyukova et al. 2009).

Most cecidomyiid SltxB sequences form a monophyletic clade sister to an unidentified bacteria isolated from *Populus* trees (Crombie et al. 2018). This clade is sister to APSE SltxB sequences, so the gene was likely originally transferred from an APSE-like ancestor (see **Figure 2** and **Figure S1**). Synteny (**Table S3**) between *C. nasturtii* and *S. mosellana sltxB* sequences show that *sltxB* was transferred to a common ancestor prior to their divergence ca. 70 mya (Dorchin et al. 2019). The gene was duplicated in *C. nasturtii* and appears in tandem copies on several scaffolds in the *S. mosellana* genome (**Table S3**). Most insect SltxB sequences form a large polytomy, consistent with a recent expansion (Whitfield & Lockhart 2007). We found several motifs involved in Gb3 binding and cytotoxicity (Bast et al. 1999) were conserved in insect and bacterial SltxB copies. Residues that contribute to cytotoxicity, including F50, A63 and G82 (Clark et al. 1996), were highly conserved between bacterial and cecidomyiid species (**Figure 3**). Phyre2 analyses show several insect SltxB sequences have retained a typical oligomer-binding fold, a SltxB structure (**Table S5**) consistent with our expectations.

Across all of the toxin genes we discovered in galling midge genomes, our phylogenetic analyses show reticulated patterns of ancestry between species. For example, in one case, APSE-like phages may not have been the ancestral donors (e.g. *lysozyme*), and, in another, HGT occurred in several insect taxa (e.g. *aip56*). Having several possible donors for HGT is not unexpected in cecidomyiids, since HGT has been reported from APSE phages (Zhao et al. 2015), fungi (Cobbs et al. 2013), and even other cecidomyiid species (Ben Amara et al. 2017). We observed that, in most cases, the phylogenies show that HGT occurred between insects and microbes associated with insects (**Figure 2, Figure S1**). The intimate associations between plant- and microbe-feeding insects, bacterial symbionts, and their phages may lead to HGT events because their DNA is in close proximity to the insect germline, which could facilitate transfer of DNA into the germline nucleus. While a simplistic model, we could find no other evidence in the literature of studies that have formally addressed if associations between insects and their symbionts facilitate HGT. We tested this ‘transfer-by-proximity’ hypothesis (that sharing similar habitats facilitates HGT) using the δ value (Borges et al. 2019), a metric for measuring the degree of phylogenetic signal for categorical traits. Phylogenetic signal is the tendency of traits from related species to resemble each other more than species drawn at random from the same tree for a phenotype (Blomberg & Garland 2002), and higher δ values correspond to higher phylogenetic signal (Borges et al. 2019). Every species on our protein phylogenies was assigned a “Lifestyle” by searching the existing literature, which included living on plants, plant roots, soil, arthropods (e.g. endosymbionts), or other habitats. We compared the δ value for the observed (“true”) trait distribution across all phylogenies with those for which the traits were shuffled randomly across the tips. We found that the δ values for the “true” trait distribution were consistently higher than the distribution of δ values when the trait was shuffled along the phylogeny (**Table S6**). Thus, there is some non-random association between habitat and transfer of genes between species. However, not all sampled tips represent HGT events, and the overabundance of vertical transfer events limited our ability to use this metric to test the hypothesis. To address this issue, we also calculated δ after removing vertically inherited tips from the phylogeny (see **Supplementary Methods**) and obtained similar results (**Table S6**). Our analysis of phylogenetic signal suggests that physical proximity of genetic material (e.g. that between eukaryotic hosts and their endosymbionts) may facilitate gene transfer. Many of the species closely related to our candidate insect genes are associated with insects, such as APSE, *Trichoplusia ni* ascovirus, or *Xenorhabdus vietnamensis*. This analysis suggests, but does not prove, that the intimate associations between these species and their insect hosts may have facilitated ancient opportunities for HGT.

Our previous knowledge of prokaryote-to-insect HGT events centered on genes involved in conferring new metabolic capabilities, particularly those that allow insects to colonize new plant hosts (Wybouw et al. 2016). HGTs of toxin-encoding genes are largely antibacterial in function, when function can be acsertained (Di Lelio et al. 2019; Hayes et al. 2020; Li et al. 2021). The galling midge HGT dataset as a whole highlights that a new functional class of proteins, toxins that antagonize eukaryotic cells, may be common among insects. Given that many of these horizontally transferred toxins (with the exception of lysozyme) target eukaryotic cellular components, the genes encoding them may have become integrated into existing immunological networks to protect cecidomyiids from attack by parasitoid wasps or other eukaryotic enemies. Notably, the cecidomyiid species sampled in our study face strong selective pressure from a wide number of parasitoid taxa (Chen et al. 1991; Abram et al. 2012; Chavalle et al. 2018). We hypothesize that *cdtB, rhs*, and *sltxB* in particular may protect developing cecidomyiid larvae and pupae from parasitoid wasps, since these three genes confer this protective function in other insects (Oliver et al. 2009; Martinez et al. 2018; McLean et al. 2018).

Our work contributes to the growing body of literature on HGT in eukaryotes, particularly of eukaryotic-targeting toxin genes. Here we also took a new approach for assessment of phylogenetic signal and found results consistent with the transfer-by-proximity hypothesis of animal HGT from microbes living in similar environments as the insects. Further sampling of genomes across Cecidomyiidae can corroborate the timing of these HGT events, in addition to revealing more about the evolution and biology of this agriculturally important insect family.

## Acknowledgments

We thank Dr. Yolanda Chen and Andrea Swan at University of Vermont for providing *Contarinia nasturtii* samples. Discussions with Dr. Iswar Hariharan and Dr. Michael Shapira motivated the phylogenetic signal analysis. Nicolas Alexandre, Dr. Carrie Malina, Dr. Jenna Ekwealor, and Dr. Jennifer Wisecaver provided valuable feedback for bioinformatic analysis and phylogenetic tree estimation. *Funding:* KIV was funded by the National Science Foundation Graduate Research Fellowship as well as grants from the Golden Gate Science into Action Fund at Golden Gate National Recreation Area and University of California - Berkeley Integrative Biology Summer Research Award. RLT was funded by the University of California - Berkeley, Berkeley Fellowship and the National Science Foundation Graduate Research Fellowship. SMA was funded by the National Institute of Health - Bridges to the Baccalaureate program, Berkeley Transfer Scholarship, and the Cal Alumni Scholarship. This work was also supported by the National Institute of General Medical Science of the National Institutes of Health award number R35GM119816 to NKW.

## Data Availability

Genomic and transcriptomic resources utilized in this text are shown in **Table S2**.

## Supplementary Methods

### Identifying horizontal gene transfer (HGT) candidates in cecidomyiids

We used HGT screening methods described in Nikoh et al. (2010), but adjusted to the scope of this study and the bioinformatic resources available. A summary of our methods is shown in **Figure S2**. To identify possible HGT candidates in the Cecidomyiidae, we ran TBLASTN on APSE proteomes against existing genomic and/or transcriptomic resources for Cecidomyiidae species (see **Table S2** for proteomic queries and Cecidomyiidae databases). These searches were conducted throughout June-July 2020. We initially retained all hits for consideration as HGT candidate Genes of Interest (or HGT-GOIs). Sequences were eliminated as HGT candidates if BLASTP searches of the predicted subject amino acid sequence (either the High-scoring Segment Pair, predicted ORF, or whole length predicted annotation) to the NCBI nr database showed the top 2+ hits were to canonical insect genes. If HGT candidates were < 50 continuous amino acids long, they were removed from consideration. Redundant hits, defined as hits where the same HGT-GOIs from different APSE strains mapped back to the same genomic coordinate, were then removed. We also removed hits encoded on scaffolds <1 kb long, as these are highly likely to be bacterial contaminants or mis-assembled regions. Additionally, we removed hits with an E-value > 0.01.

### Quality Control (QC) for HGT candidates

We considered strong evidence of HGT if the gene of interest (henceforth referred to as ‘GOI’) met 2/5 of the following criteria:

i. Non-anomalous read depth via BWA analysis (**Supplementary File 1**)
ii. The GOI was on scaffolds with other *bona fide* eukaryotic genes (**Supplementary File 1**)
iii. The GOI is syntenic in two or more species (**Table S3**)
iv. The GOI is transcribed in dT-enriched transcriptomes (**Table S2**)
v. The GOI is predicted to have introns (**Supplementary File 1**)

In the case of *Contarinia nasturtii*, we validated the HGTs with PCR and bi-directional Sanger sequencing (see **Supplementary Methods** and **Table S4**) of genomic DNA isolated from larvae and adults of this midge species. In cases where the distance between the GOI and a proximal gene was <2000 bp, we amplified regions that included other *bona fide* eukaryotic genes. A summary of our QC methods for each species is shown in **Supplementary File 1**. Our final list of HGT candidates is shown in **Table 1**.

To determine if multiple HGT-GOIs were actually duplicates or a consequence of mis-assembly, we compared the scaffolds of gene duplicates using progressiveMauve (Darling et al. 2010). If there was >90% nucleotide identity between scaffolds, we considered those mis-assembly artifacts. For tandem duplicates, we further used BWA to analyze read depth at each paralog to determine if one or all paralogs had anomalous read depth that could signal mis-assembly. Additionally, if encoded genes were <10% of the size of the canonical, functional protein, they were discarded as candidates.

A simplified workflow to identify functional HGT candidates is shown in **Fig S2**.

### Burrows Wheeler Alignment (BWA) analysis

We aligned Illumina reads (see **Table S2** for SRA accessions) to the genome via BWA (Li & Durbin 2009) to search for unusual coverage depth relative to neighboring genes, which can be due to contamination (Koutsovoulos et al. 2016). Read quality and trimming were assessed with FastQC (Andrews 2010), which showed high per base sequence quality, low per base N content, and low adapter content. The read alignment was visualized and assessed in the software package Geneious v. 11.1.5 (https://www.geneious.com). Since the majority of the genes were encoded on scaffolds encoding other *bona fide* eukaryotic genes, we included the read depth of all candidate scaffolds, per species, in a Grubbs’ test and removed scaffolds with read depth outliers. Following this, we did the same with the loci containing the horizontally transferred genes (HTGs). The results show there are no coverage abnormalities, indicating these candidate HTGs are not due to assembly artifacts or microbial contamination.

### Transcription analysis

We submitted the GOI (+/- up to 20 kb up and downstream) as a blastn query to representative polyA-enriched transcriptomes. These representative transcriptomes are shown in **Table S2**. The top hits (≤ 5000) were extracted and mapped back to the region using Geneious RNA Mapper (Sensitivity: Highest Sensitivity / Slow; Span annotated introns). We report the mean read depth and standard deviation across the GOI. Limitations of the transcriptional analysis are that characterizations of any given transcriptome are contingent on tissue type and life stage, and as such, the expression patterns of one transcriptome may not reflect the importance of the gene in a species.

### Synteny analyses

Possibly due to the high divergence between sequenced species (e.g. *C. nasturtii* and *S. mosellana* are estimated to have split ∼70 mya [Dorchin et al. 2019]), macro-syntenic analyses using progressiveMauve (Darling et al. 2010) and CoGe SynMap (Lyons et al. 2008) were not fruitful. Instead, we employed a qualitative micro-syntenic approach. In annotated genomes, we extracted the protein sequences of genes up and downstream of the GOI and indicated their position with -*n* or +*n* (for example, a positionality of −3 indicates the gene is located three genes upstream of the GOI). These sequences were then submitted as TBLASTN queries (Altschul et al. 1997) to the representative genomes. The scaffolds of top hits were then extracted; if there were no hits, we indicate “NA” in the cell. We considered there to be some evidence of synteny if one or more genes proximal to the GOI were located on the same scaffold within a species. Results are shown in **Table S3**.

### gDNA extraction and PCR conditions

Ethanol-preserved samples of *C. nasturtii* larvae and adult males and females from a lab-reared colony were provided to us by Andrea Swan and Dr. Yolanda Chen at University of Vermont. Ethanol-preserved specimens were rehydrated in sterile water and allowed to dry on a Kimwipe prior to DNA extraction. Rehydrated specimens were homogenized with a bead-beater for 2 minutes at 30Hz. DNA was extracted from the homogenized samples using a DNEasy Kit (Qiagen) with an overnight Proteinase K digestion at 55°C. gDNA samples contained 1-5 larvae or a single adult midge.

We designed PCR primers to capture the HGT-GOI and, if the nearest predicted gene was <2kb distant, a neighboring *bona fide* eukaryotic gene. PCR primers were designed using Geneious. PCR reaction mixes were composed of: 7.5µl Failsafe Premix E (Epicentre), 4.2µl nuclease-free water, 1.2µl each of F and R primers (IDT), 1µl template DNA, and 0.12µl *Taq* polymerase (New England Biolabs). Thermocycler settings were: 5 m at 95°C and 35 cycles of 95°C for 30 s, Ta for 30 s, and 68°C for 1-2.5 m depending on amplicon size (see **Table S4**), followed by a final 5 m extension at 68°C. 1% agarose 1X TBE gels were prepared with Apex Agarose in 1X TBE buffer with 1 µL SYBR™ Safe staining gel per 10 mL of gel solution. 4 µL of PCR product was mixed with 1 µL ThermoScientific 6X Loading Dye. 1Kb Plus DNA ladder (Invitrogen) was included as a molecular marker. PCR product was run on gels using the Owl™ EasyCast™ B1 Mini Gel Electrophoresis System rigs for 30 minutes at 120V. Gels were visualized using AlphaImager™ Gel Imaging System (Alpha Innotech). Primer sequences, the region captured by the amplicon, melting temperature, extension times, and the expected amplicon length are detailed in **Table S4** along with gel images. All PCR products were Sanger sequenced in both directions at the UC Berkeley DNA Sequencing facility.

### Species phylogeny and ancestral state reconstruction

Nucleotide sequences for *co1, cad, ef1a*, and *28S* were retrieved from GenBank for each of the five species included in the species phylogeny (**Table S7**). *Bibio marci* (Diptera: Bibionidae) was included as an outgroup to the Cecidomyiidae family, consistent with phylogenies previously generated for the family (Sikora et al. 2019). Each gene was aligned individually using the default settings on the MAFFT v. 7 web server (Katoh et al. 2019). Individual gene alignments were inspected and manually trimmed before concatenation. The final alignment consisted of five species and a total of 3135 nucleotide sites. Total sequence lengths for each gene were as follows: *co1* (542 nt), *cad* (1439 nt), *ef1a* (725 nt), *28S* (429 nt). The concatenated alignment was uploaded to CIPRES web portal for maximum likelihood (ML) tree construction. A ML tree was generated using RAxML-HPC2 on XSEDE using default settings (Miller et al. 2010; Stamatakis 2014). The ML species tree is shown (log likelihood = −10311.662040) with bootstrap values at each node (*n* = 1000 bootstraps) (**Figure 1**).

Due to the low number of taxa on our tree, maximum likelihood approaches to timing HGT events were uninformative. We used maximum parsimony (MP) to infer the timing of each HGT event by incorporating data from synteny analyses and protein phylogenies. Briefly, we assumed a single acquisition of the HTG in the common ancestor if there was evidence of shared synteny among the taxa in which the HTG was found (**Table S3**). In the absence of synteny data, we examined the protein phylogenies to determine timing of HGT events (**Figure S1a**). We interpret monophyly of cecidomyiid protein sequences as a single acquisition event in a common ancestor under a MP model. Acquisition events that are only supported by protein phylogeny data are indicated on the species tree with dashed ticks (**Figure 1**).

### Protein phylogeny construction

Representative toxin sequences were queried against the NCBI refseq protein database on 11/20/2020 using BLASTP (Altschul et al. 1997) with a maximum of 500 top hits per query (see below for a list of query sequences used per toxin). Top hits were extracted for each sequence, and redundant sequences were removed with cd-hit (Li & Godzik 2006; Huang et al. 2010) with a 0.8 similarity cutoff, unless they were genes specifically identified in this manuscript.

Sequences were aligned with MAFFT v. 7.312 using the E-INS-I strategy and the BLOSUM62 amino acid scoring matrices (Katoh & Standley 2013). Sequences were trimmed to include only the conserved protein domains (i.e., domains in which <50% of the sequences had gaps). After trimming, sequences were re-aligned with the earlier MAFFT settings. Gene topologies were inferred using maximum likelihood as implemented in W-IQ-TREE (http://iqtree.cibiv.univie.ac.at/) (Nguyen et al. 2015; Trifinopoulos et al. 2016) using the best-fit model as assessed by BIC in ModelFinder (Kalyaanamoorthy et al. 2017). The resultant consensus tree was constructed from 1000 ultrafast-bootstrapped trees (Hoang et al. 2018). Nodes with <50% bootstrap support were collapsed to polytomies using the di2multi function in ape v5.4 (Paradis & Schliep 2019).

Life history of represented sequences in each phylogeny were determined by searching the literature (see **Supplementary File 2**) while Phylum information was taken from NCBI Taxonomy (Sayers et al. 2019; Schoch et al. 2020). Life history and phylum information were used to annotate the phylogeny. Phylogenies were visualized and annotated using ggtree v. 2.5.0.991 (Yu et al. 2017; Yu 2020).

### Specifics of each phylogeny are shown below

1. **Aip56 queries:** Queries: *Drosophila bipectinata* (XP_017099943.1, AIP56 domain), *C. nasturtii* (XP_031636937.1, XP_031641113.1), *M. destructor* (this manuscript), *S. mosellana* (this manuscript), APSE1 (NP_050970.1), APSE4 (ACJ10093.1), APSE5 (ACJ10079.1). Total tips: 90. Model: LG+G4. LogL= −40095.6183, BIC=81322.3145.
2. **CdtB**. Queries: Candidatus *Hamiltonella defensa* (XP_016857353.1), *C. nasturtii* (XP_031641203), *D. ananassae* (XP_014760894.1), *D. biarmipes* (XP_016950904.1), *Myzus persicae* (XP_022165116.1), and *Scaptomyza flava* (QDF82162.1). Total tips: 76. Model: LG+F+I+G4. LogL = −27554.2607, BIC = 56112.4328.
3. **Lysozyme queries:** Queries: APSE2 (YP_002308525.1), *C. nasturtii* (XP_031638744.1), *M. destructor* (this manuscript). To make the tree clearer, we removed one large and highly divergent clade from the phylogeny which included *Hamiltonella defensa* and APSE-2 phage sequences. Total tips: 172. Model: WAG_I_G4. LogL=-43487.1457, BIC = 90213.0519.
4. **SltxB queries:** Queries: APSE1 (NP_050968.1), APSE5 (ACJ10077.1), *C*.*nasturtii* (XP_031619577.1, XP_031619578), *Burkholderia ambifaria* (WP_175804727.1), and *S. mosellana* (copies identified in this manuscript). Total tips: 28. Model: VT+G4, LogL=- 3729.8063, BIC=7016.7700.
5. **RHS queries:** Queries: Bacteriophage APSE3 (CAB3775397.1), *Candidatus* Hamiltonella defensa (ATW32053.1), *S. mosellana* (copies identified in this manuscript), and *M. destructor* (Mdes011034). Total tips: 188. Model: WAG+F+I+G4. LogL = - 312544.0798, BIC = 628028.7246.

### Measuring phylogenetic signal

For all species in a phylogeny, we assigned a “Lifestyle” trait that fell under *Arthropod, Plant, Nematode, Plant Root, Soil*, or *Other*, assignments that were meant to generally describe the lifestyle of the species. *Other* included mammalian pathogens, free-living oceanic bacteria, synthetic constructs, or other lifestyles that did not fall under the named categories. We searched the literature to determine these assignations, and citations for all species that are not *Other* are shown in **Supplementary File 2**.

We utilized Borges’ δ value to evaluate phylogenetic signal of the species’ lifestyle traits (Borges et al. 2019). The value of δ can be any positive real number. The higher the number, the higher the phylogenetic signal (Borges et al. 2019). This can be compared with the δ value of the same tree with randomized or shuffled traits to assess significance. To determine whether to “shuffle” traits (i.e. re-arrange the traits) or randomly assign traits, we piloted this analysis with shuffled and randomized trait sets, using lambda = 0.1, se=0.5, sim = 10,000, thin=10 and burn=100 in R (R Core Team 2017). We found that the shuffled trait set has a higher δ value, and as such is a more conservative method that we continued to implement.

We calculated the δ value using lambda = 0.1, se=0.5, sim = 10,000, thin=10 and burn=100 in R (R Core Team 2017). The originally calculated phylogenies were used, with one modification. We did not utilize the di2multi() function in ape (Paradis & Schliep 2019) which was implemented in our original phylogenies in order to be compatible with the δ calculation. To determine whether the realized δ value is statistically significant, we randomized the trait *n*=100 times along the phylogeny and calculated δ for each shuffling using the replicate() function in R (R Core Team 2017). The real value was compared to the randomized distribution of δ values. P-value was calculated as the number of simulations in which the shuffled δ is higher than the realized δ, a strategy utilized in several recent studies (Gruson et al.; Pinna et al. 2020; Ronget et al. 2020). δ values for the actual tree and trait assignments are reported in **Table S6**.

One major caveat is that not all sampled tips represent HGT events, and the overabundance of vertical transfer events may limit our ability to use this metric to test the hypothesis that similar lifestyles between organisms facilitates HGT. To improve the robustness of our conclusions, we removed vertically inherited tips from our phylogenies. To analyze our phylogenies *without* vertical descendance, we used the phylogenies calculated above. Then, we used the drop.tip() function in ape (interactive=TRUE) to manually remove one tip per sister taxa, *if* the two sister taxa were both from the same genus. One exception is in the case of the SltxB phylogeny, where we removed multiple cecidomyiid tips due to strong evidence of vertical descendance, as described in the manuscript. This process was repeated iteratively until the final trimmed tree had no sister taxa from the same genus. We show a representative example of the two trees, with vertical descendance and without, side-by-side in **Figure S3**.We then calculated the real and shuffled δ values as described above on the pruned tree (shown in **Table S6**).

We acknowledge there are still flaws in the above designs. First, our protocol may not eliminate all instances of vertical transfer, as vertical descendance will often link members of different genera. Furthermore, our protocol may also erroneously remove some instances of horizontal gene transfer, as we consider there may theoretically be transfer of genes *within* a genus. Additionally, the oversimplification of “Lifestyles” could lead to artificially high δ values.

### Structural analysis

To model and predict protein structure and function for representative proteins, we used the Phyre2 web portal (Kelley et al. 2015) using the “Normal” modeling mode (**Table S5**).

To determine at the residue level whether or not vital catalytic residues are preserved in disparate lineages, we used the MAFFT aligner as described above with representative sequences (Katoh & Standley 2013). The representative sequences shown in **Figure 3** are indicated in **Table S8**. Note that due to the low conservation between Aip56 insect and characterized sequences, we did not show this alignment. Alignments shown in **Figure 3** were visualized using BoxShade (https://embnet.vital-it.ch/software/BOX_form.html).

## Supplementary Text

### Domestication of various bacterial toxins following horizontal gene transfer from prokaryotes to eukaryotes

The presence of eukaryotic transcriptional motifs in putative HTGs may indicate domestication of a gene of prokaryotic origin in its novel eukaryotic context. Here, we analyze HGT sequences for motifs related to eukaryotic transcription, largely following methods described in Verster et al. (2019). Briefly, we analyzed the regions flanking our candidate HTGs for core promoter elements identified by transcription initiation factors TFIID and TFIIB (summarized in (Thomas & Chiang 2006)), alternative transcription initiation elements such as the GC box (Blake et al. 1990) or CAAT box (Graves et al. 1986; Raymondjean et al. 1988), and transcription termination elements such as polyadenylation signals, cleavage sites (CA), and upstream and downstream sequence elements (summarized in (Proudfoot 2011)). We also searched the sequences for the Shine-Dalgarno sequence, a motif essential for bacterial ribosome binding (Shine & Dalgarno 1974), which can indicate that our HTG may be a bacterial contaminant, as well as motifs for eukaryotic translational start sites, like Kozak sequences (Cavener 1987). This list is not exhaustive, nor will every element described above necessarily be found in all eukaryotic genes (Kutach & Kadonaga 2000).

We did not analyze candidate HTGs from *S. mosellana* since the genome was unannotated, making it difficult to accurately predict gene boundaries. Additionally, we did not analyze the horizontally transferred lysozyme copies, since phylogenetic analyses indicate these were transferred from a eukaryotic donor (see **Main Tex**t, **Figure 3** and **Figure S1**).

#### C. nasturtii

**Legend:**

- Predicted exons are highlighted in blue and predicted introns are highlighted in yellow. Exon/intron boundaries for *C. nasturtii* are taken from the GenBank assembly annotations.
- Coding sequences are indicated in bold text. 5’ and 3’ UTRs are therefore designated by unbolded text highlighted in blue.
- Poly(A) signals or cleavage sites are underlined. Upstream and downstream sequence elements are italicized.
- Intergenic regions (between *sltxB* copies) are designated by lowercase text.
- TATA box motifs are designated in orange. Initiator sequences are highlighted in white text when found within annotated mRNAs, or in blue text if found outside the gene boundaries. Kozak sequences are highlighted in green.

*cdtB* on NW_022197544.1:

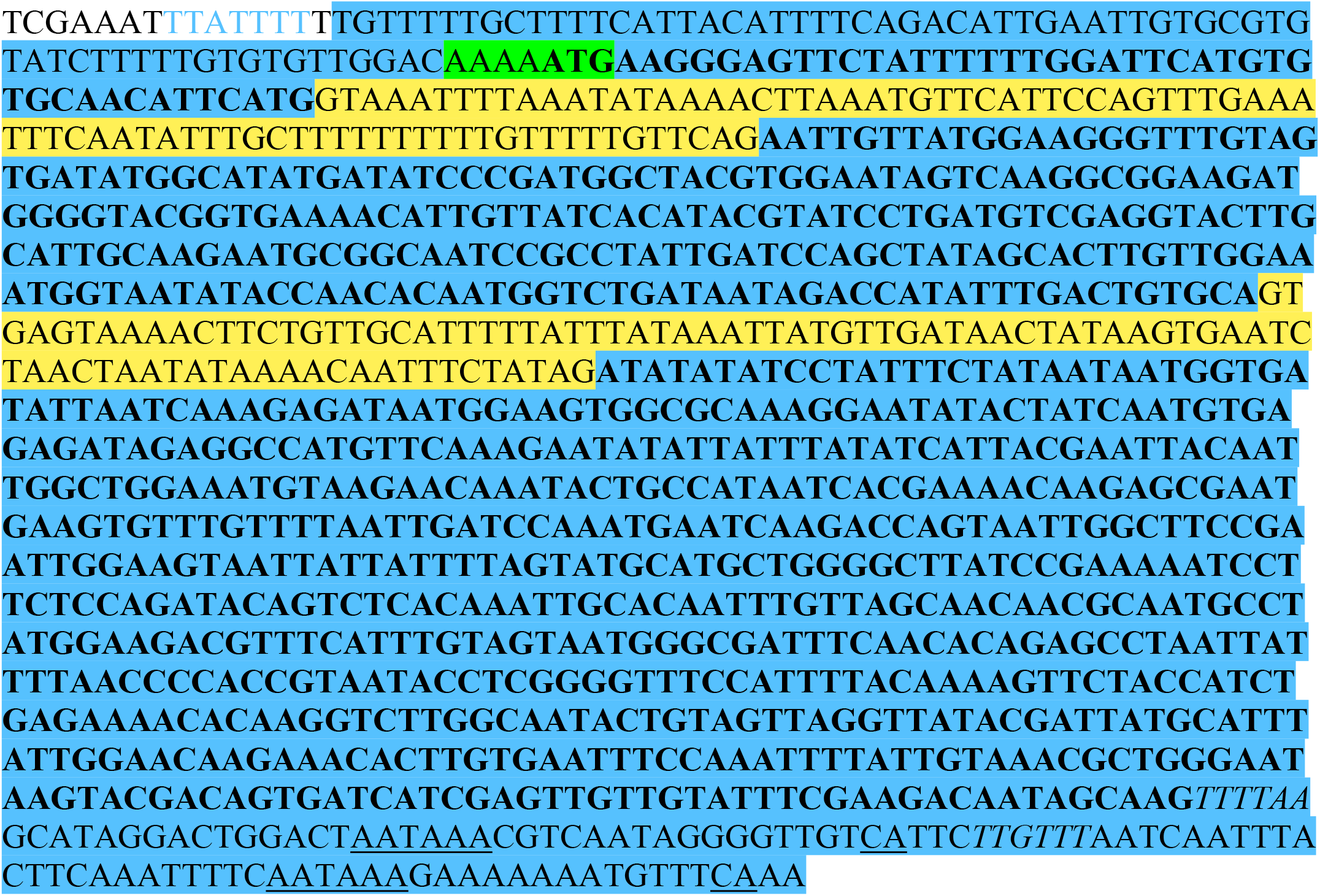

*cdtB* on NW_022203704.1:

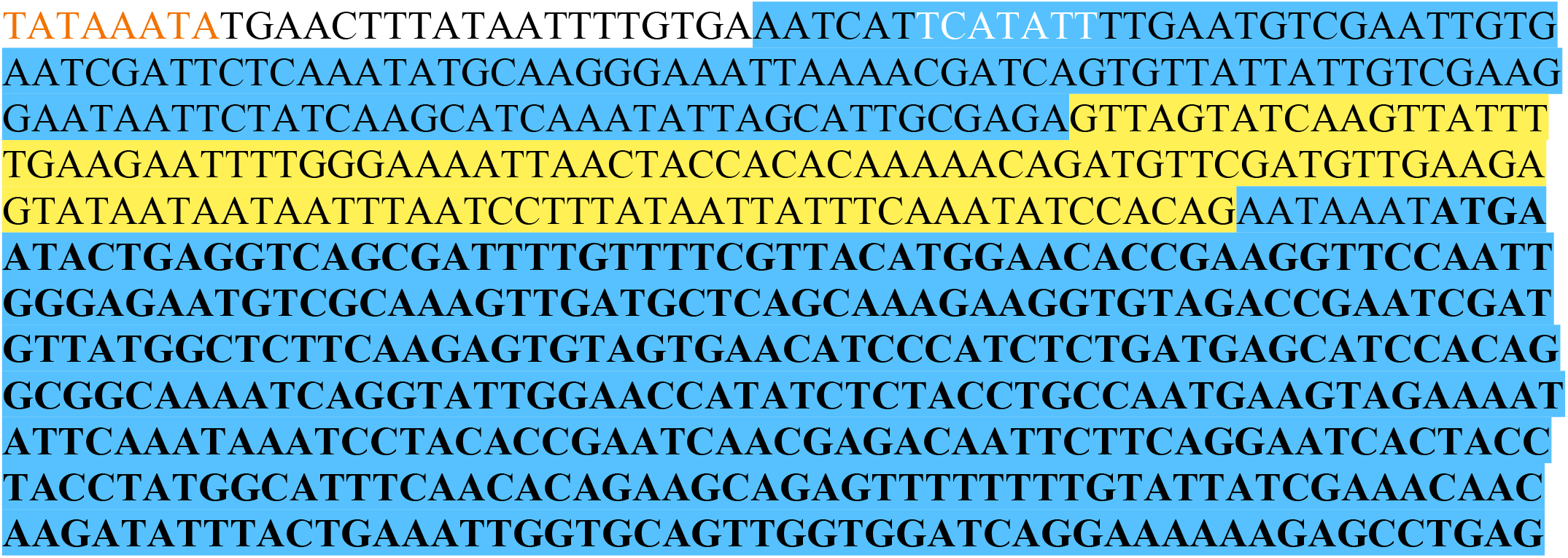
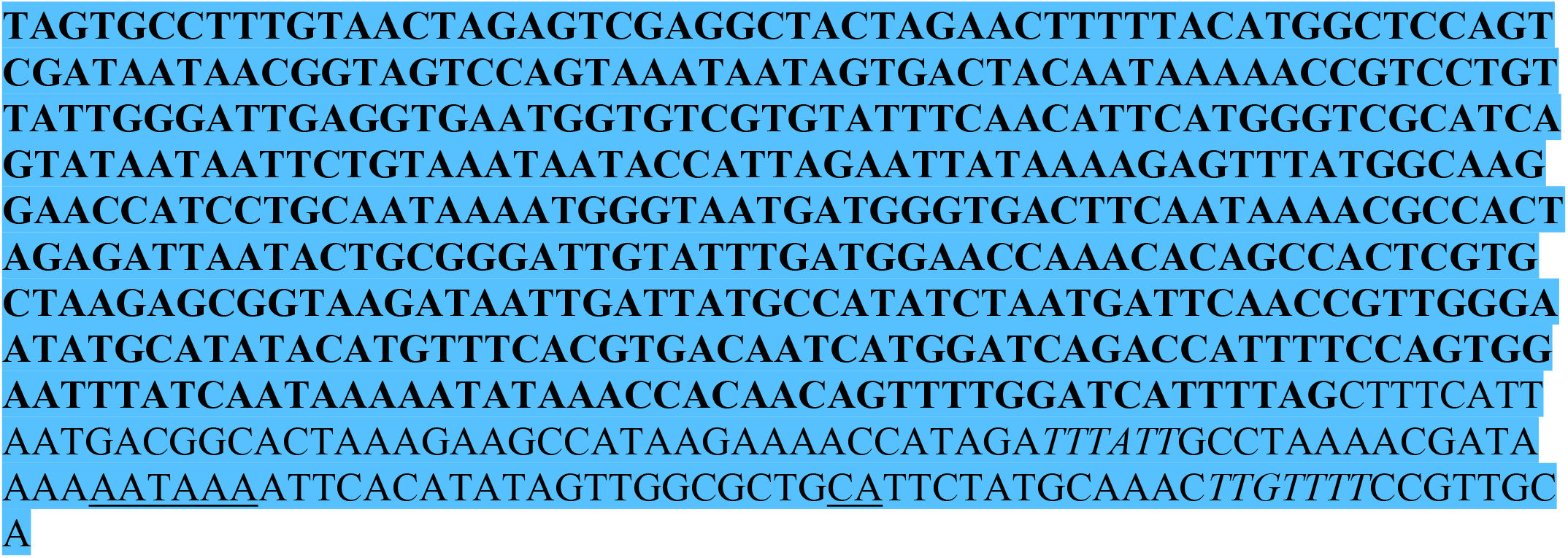

*aip56* on NW*_*022197544.1:

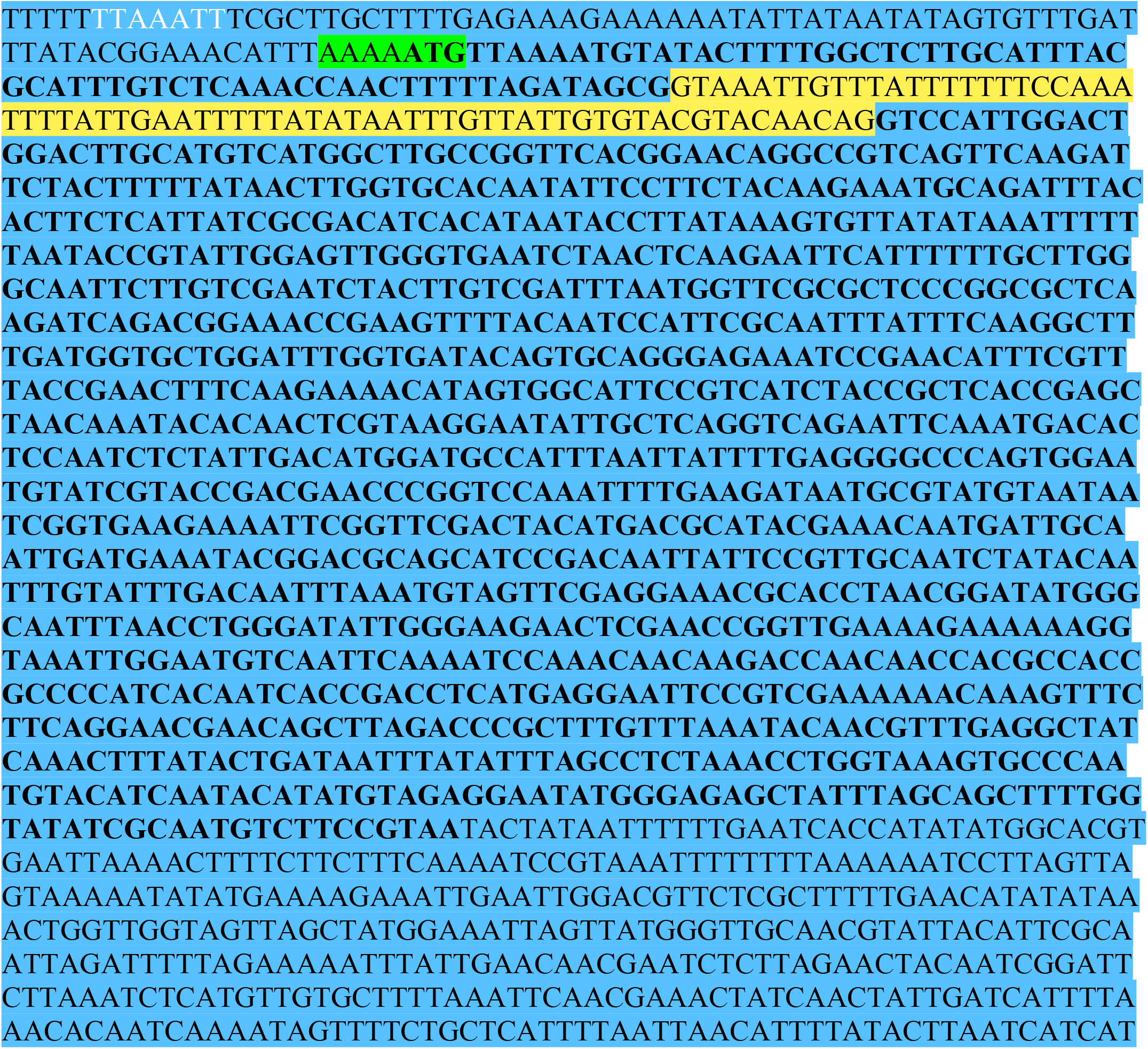
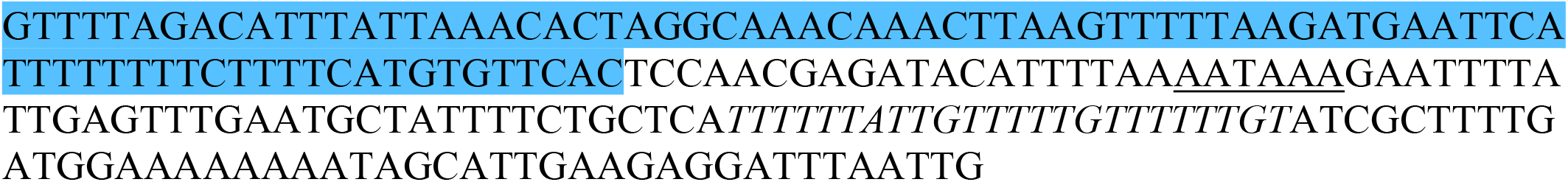

*aip56* on NW_022200251.1:

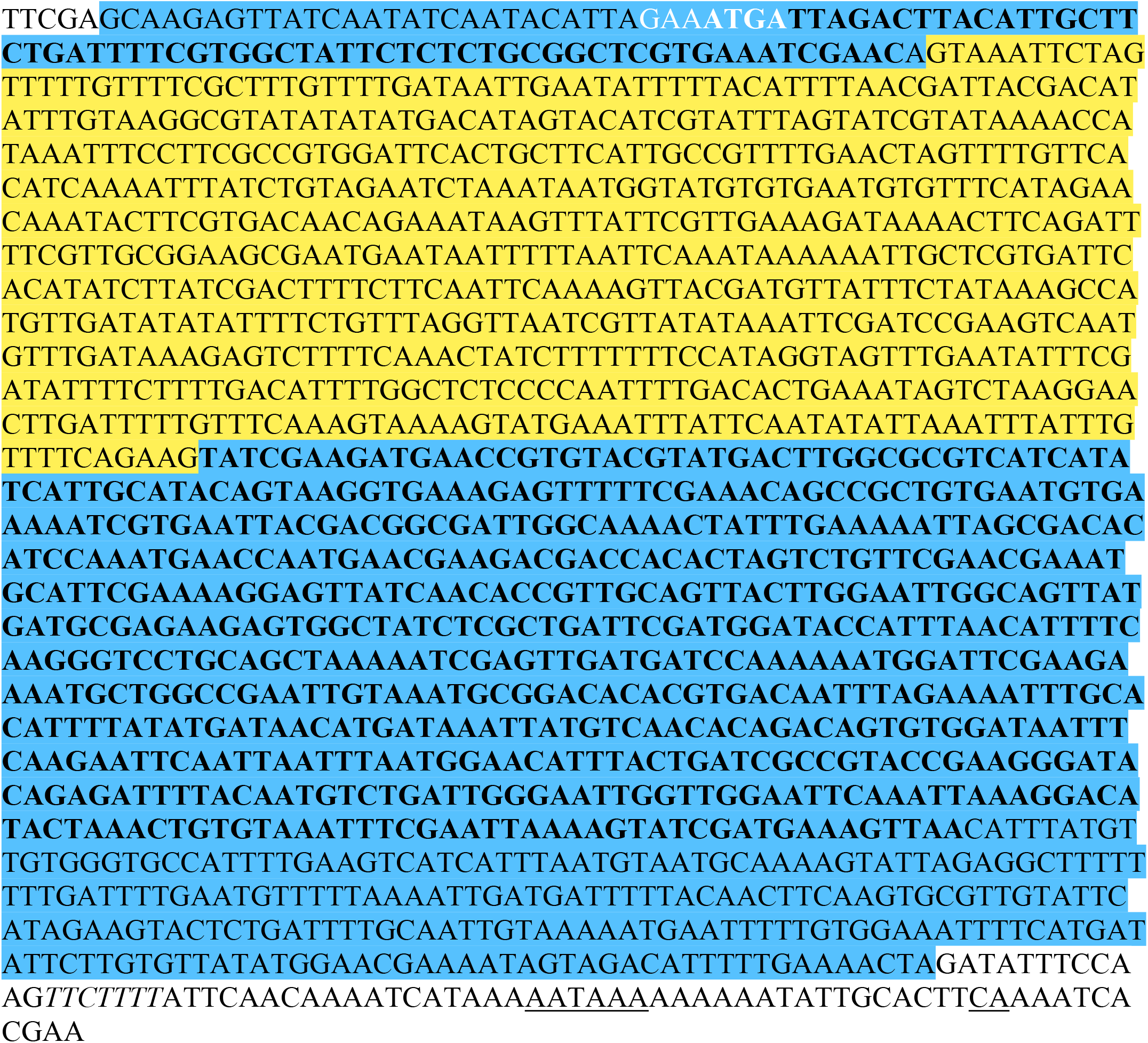

*sltxB* on NW_022197768.1:

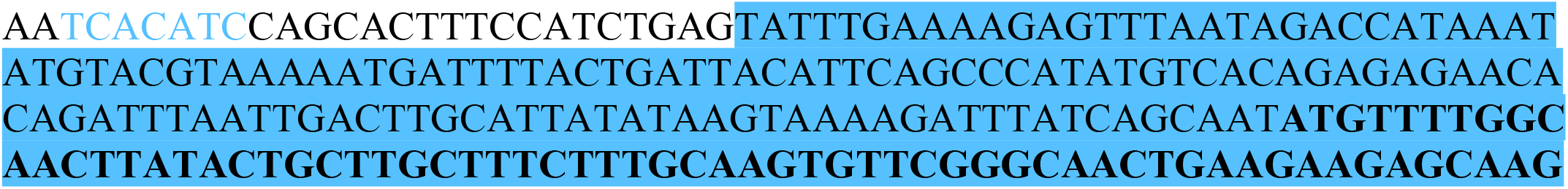
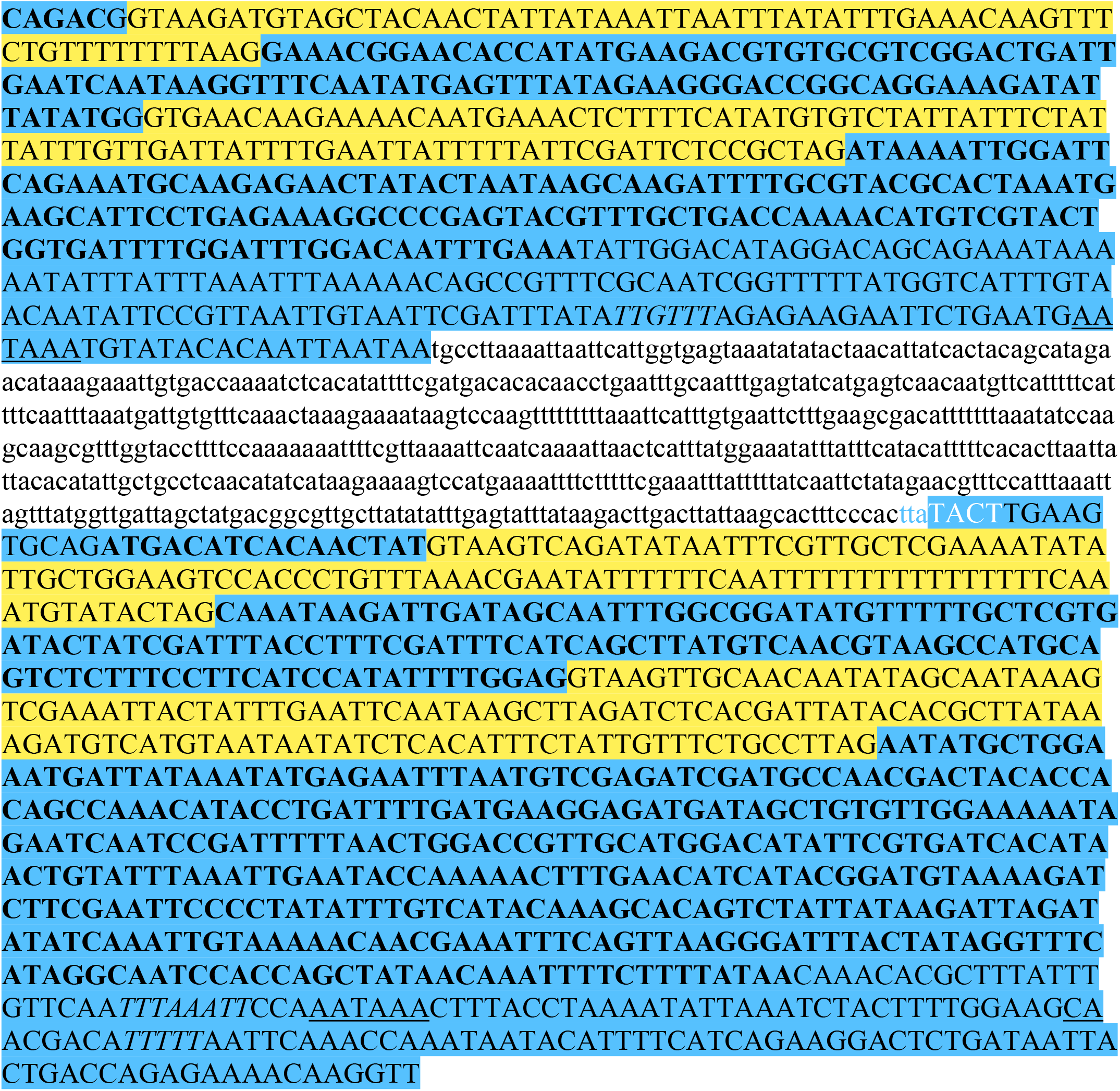

#### M. destructor

**Legend:**

- Predicted exons are highlighted in blue and predicted introns are highlighted in yellow. Exon/intron boundaries for *M. destructor* are taken from Ensembl.
- Coding sequences are indicated in bold text. 5’ and 3’ UTRs are therefore designated by unbolded text highlighted in blue. The sequences for unannotated HGT candidates are highlighted in orange, with putative in-frame start and stop codons indicated in bold orange text.
- Poly(A) signals or cleavage sites are underlined. Upstream and downstream sequence elements are italicized.
- Initiator sequences are highlighted in white text when found within annotated mRNAs, or in blue text if found outside the gene boundaries.

RHS on AEGA01002600.1:

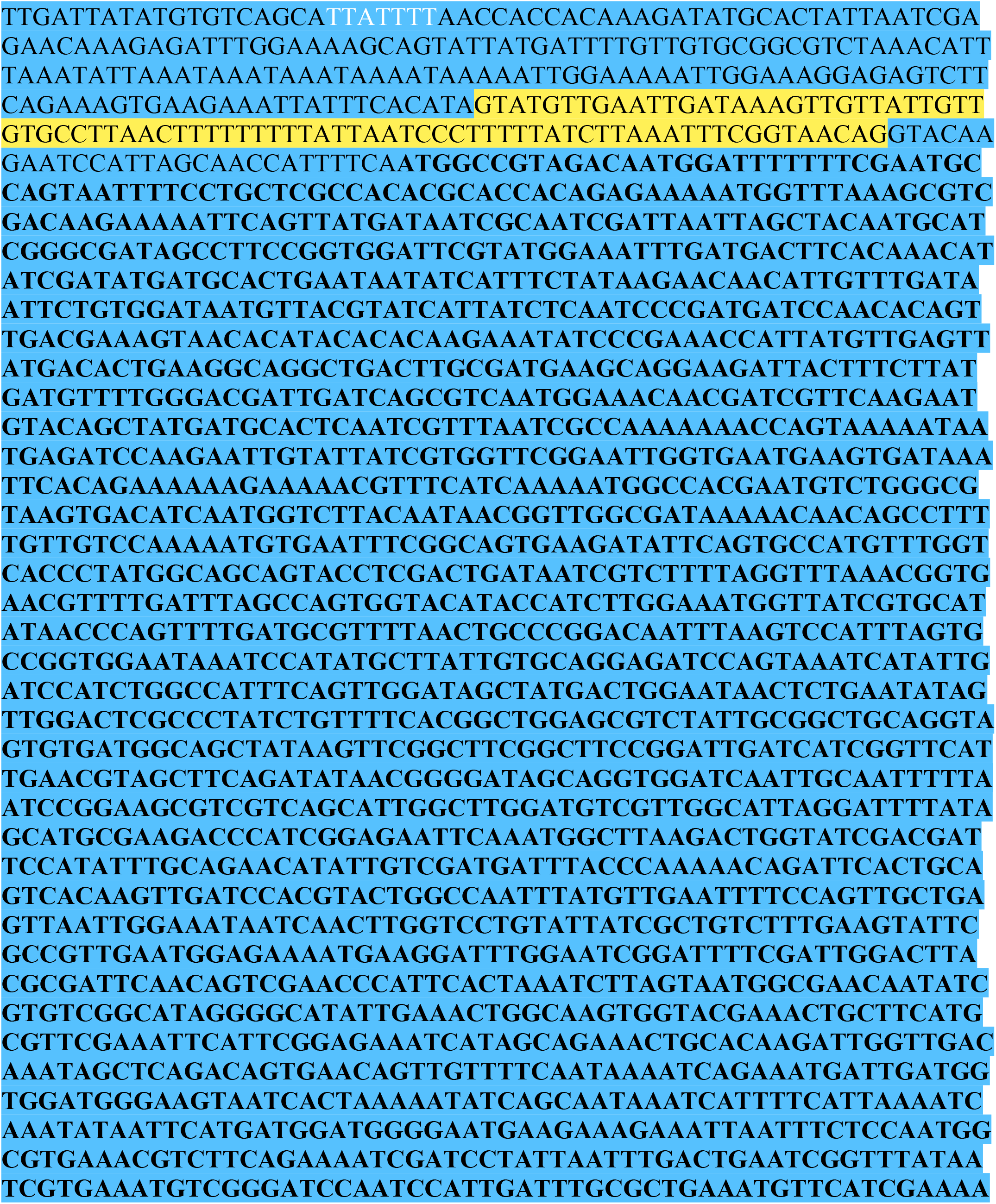
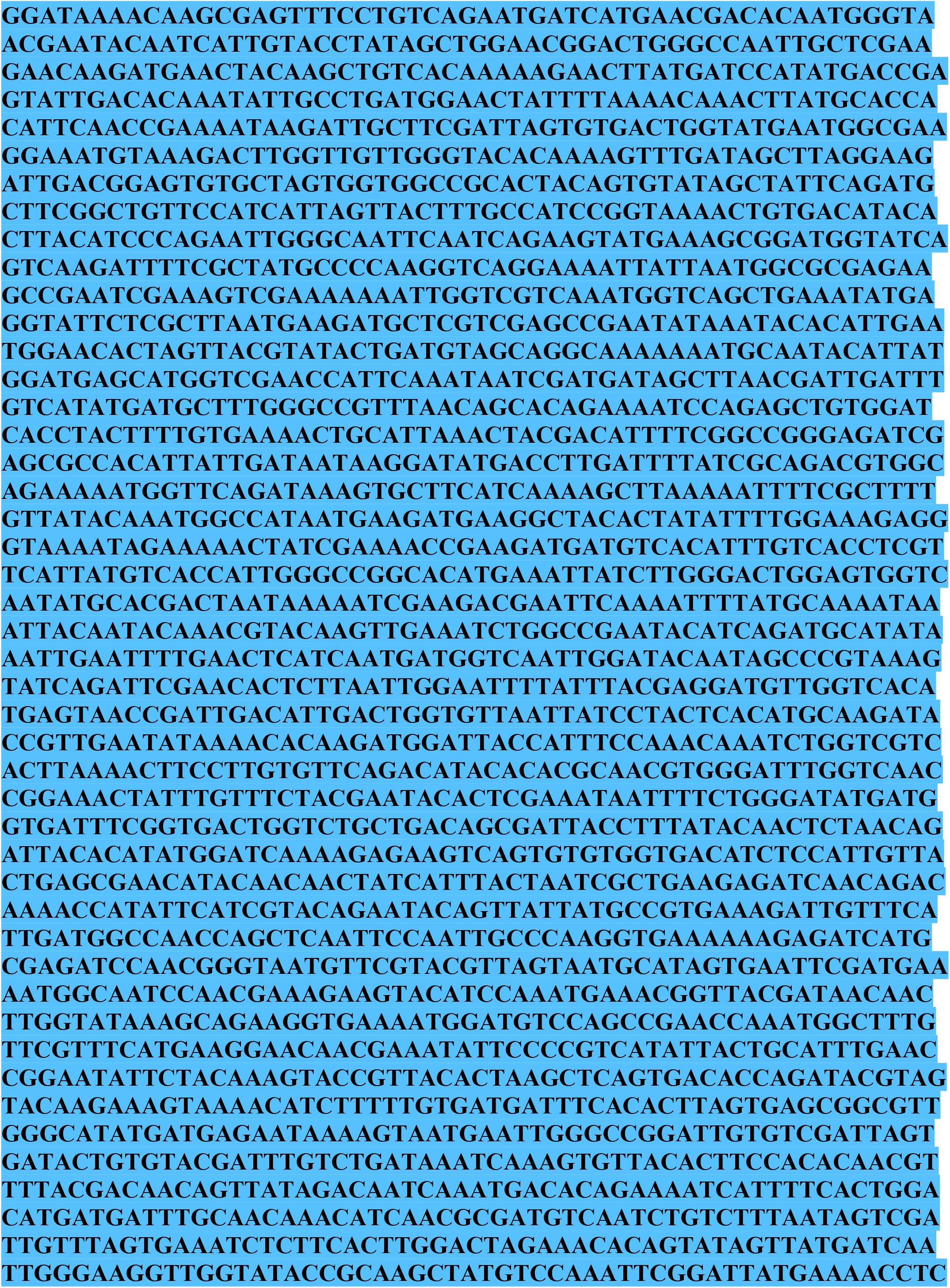
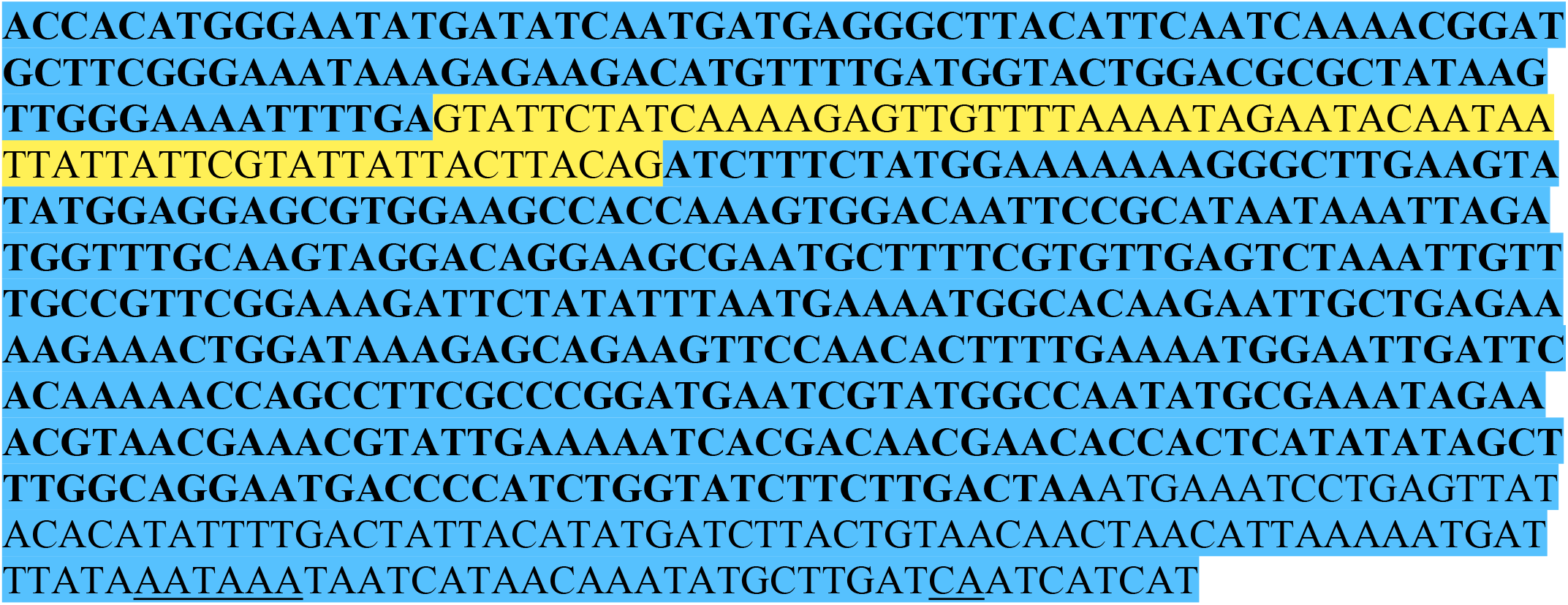

Aip56 on AEGA01003780.1:

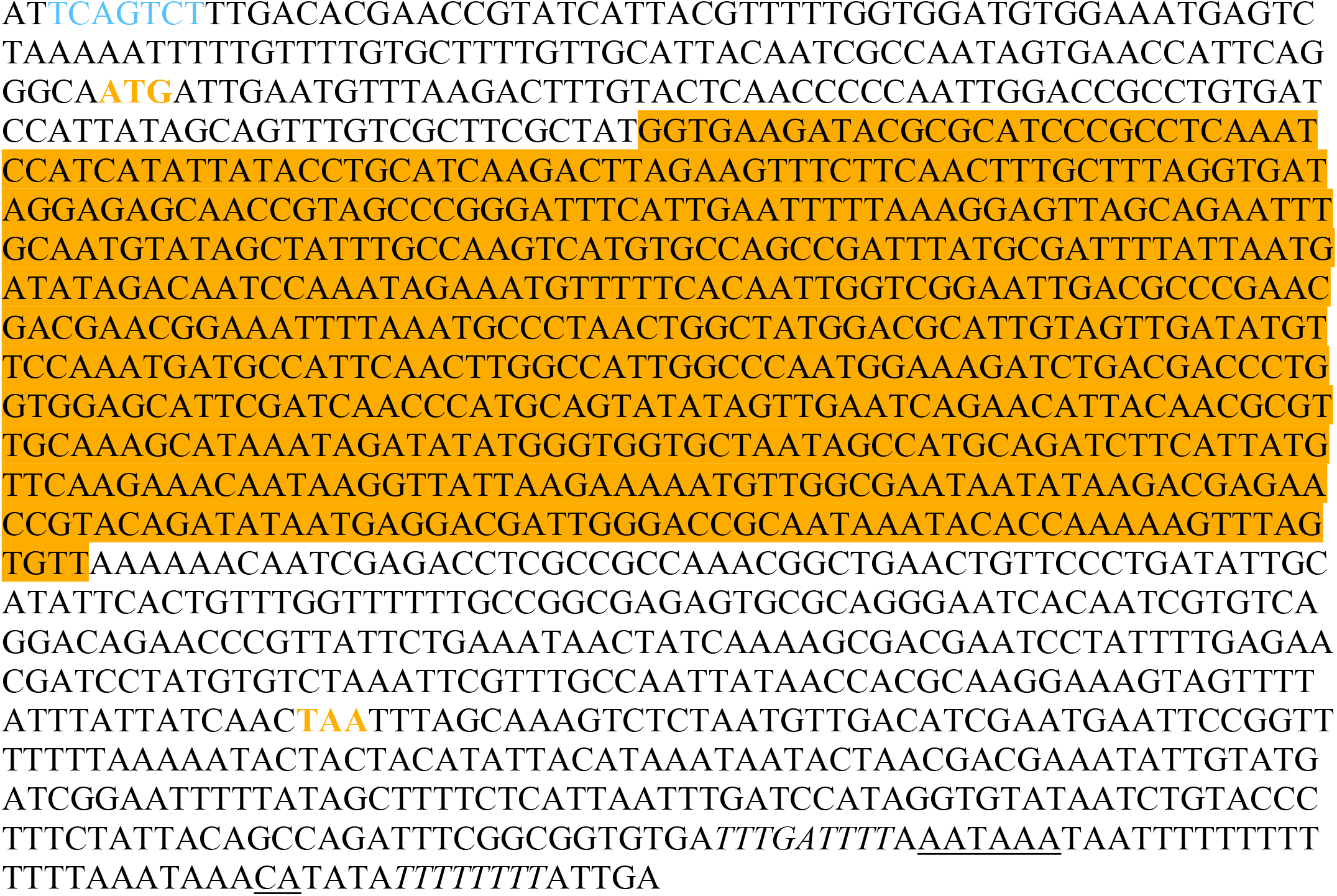

**Table S1.**
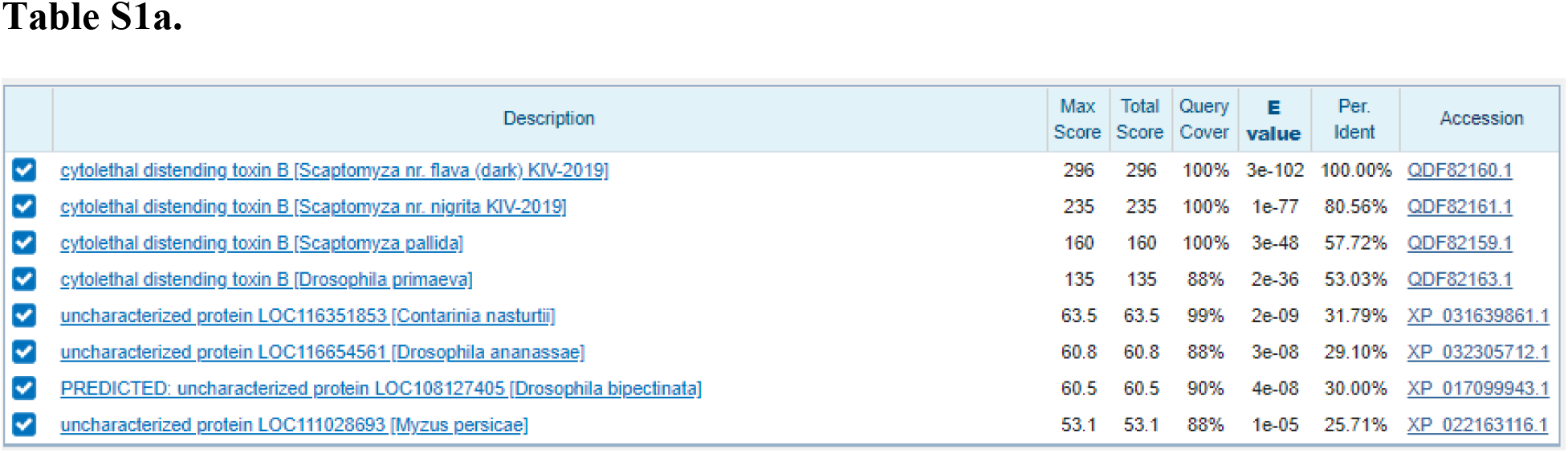
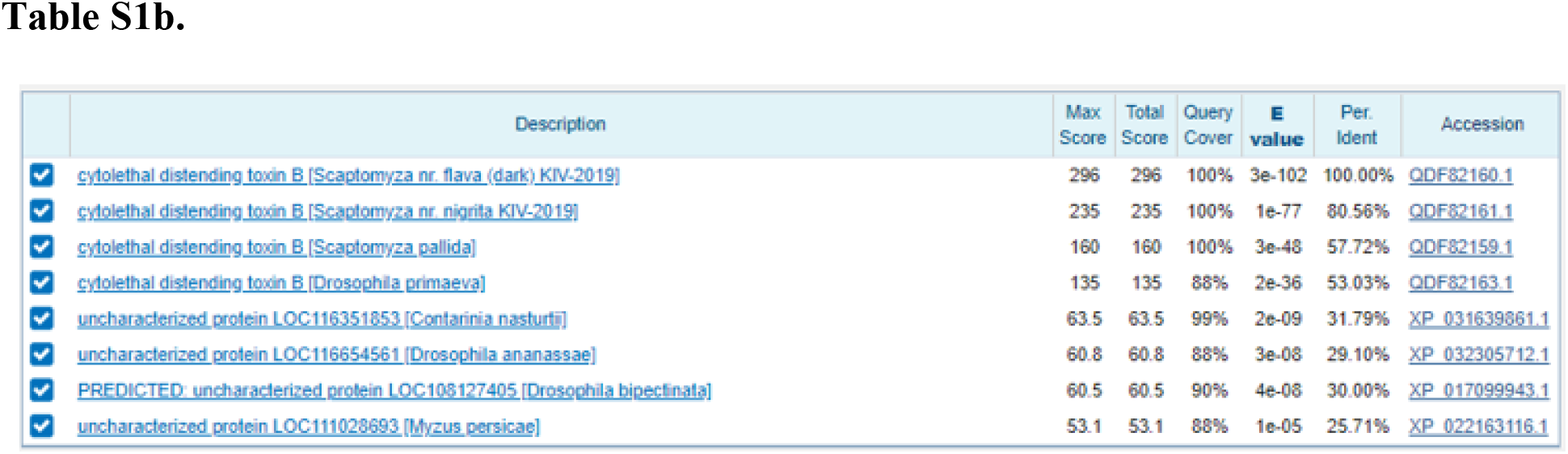
CdtB sequence QDF82160 from *Scaptomyza* nr. *flava* was used as a BLASTP query to the NCBI GenBank: Eukarya (taxid: 2759) database on 9/28/2020. These results show that *C. nasturtii* was a hit (**Table S1a)**. Another search with this newly discovered sequence, XP_031639861.1, as query shows that there are two possible *cdtB* copies in the *C. nasturtii* genome (**Table S1b**).

**Table S2.**
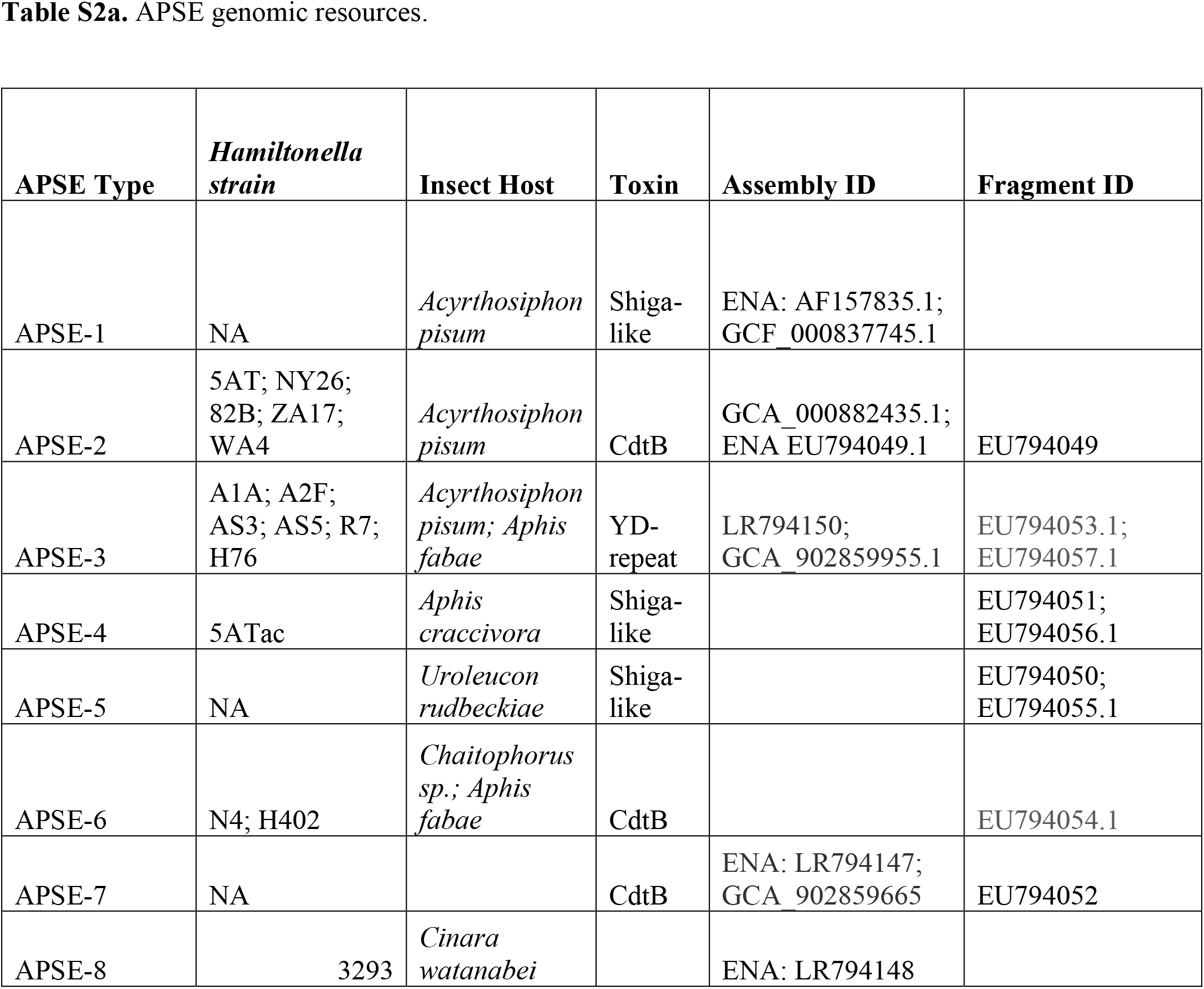
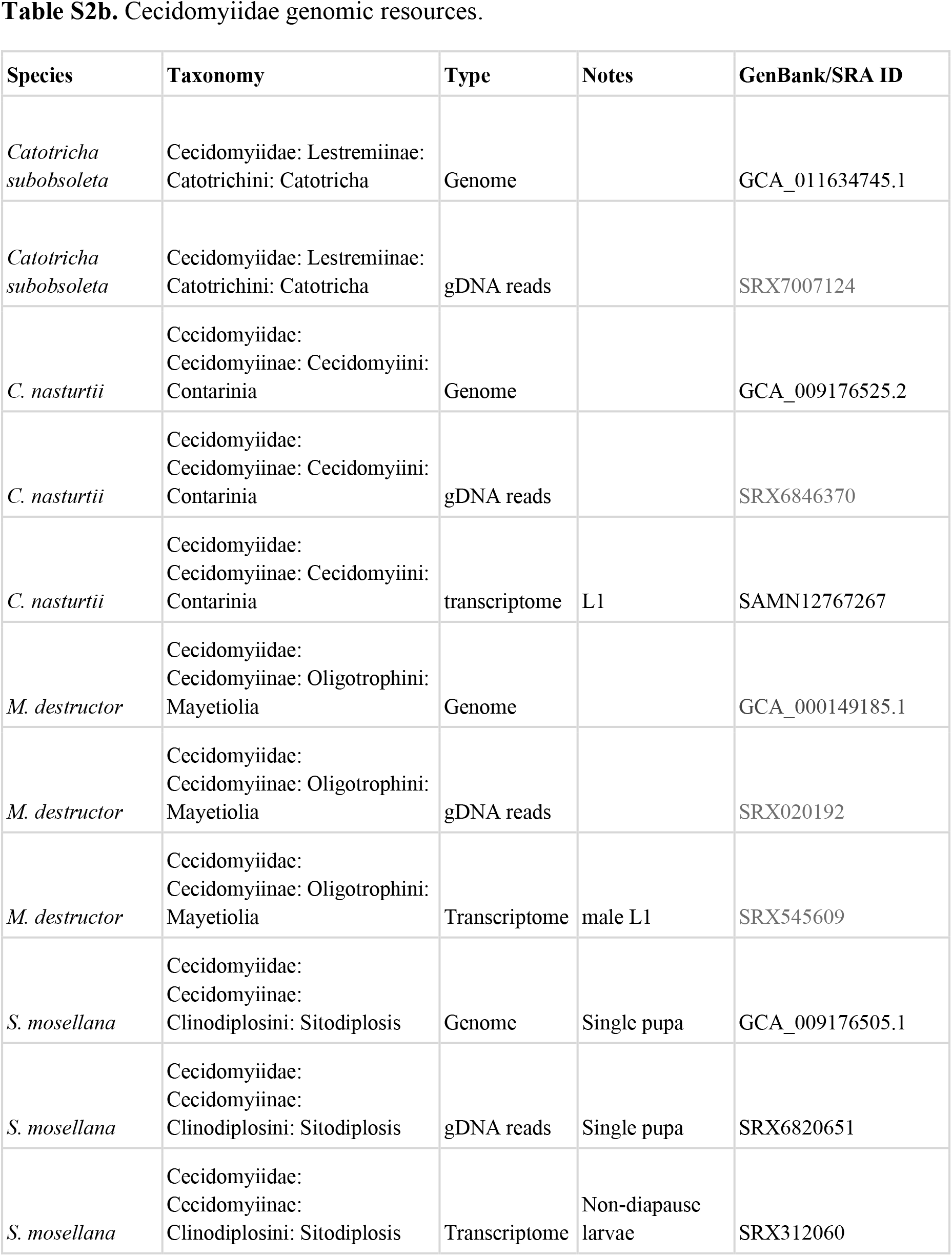
Genomic and transcriptomic resources utilized in the text. **Table S2a** shows APSE genomes whose proteomes were used as queries against cecidomyiid genomes. **Table S2b** includes interrogated cecidomyiid genome and transcriptome information.

**Table S3.**
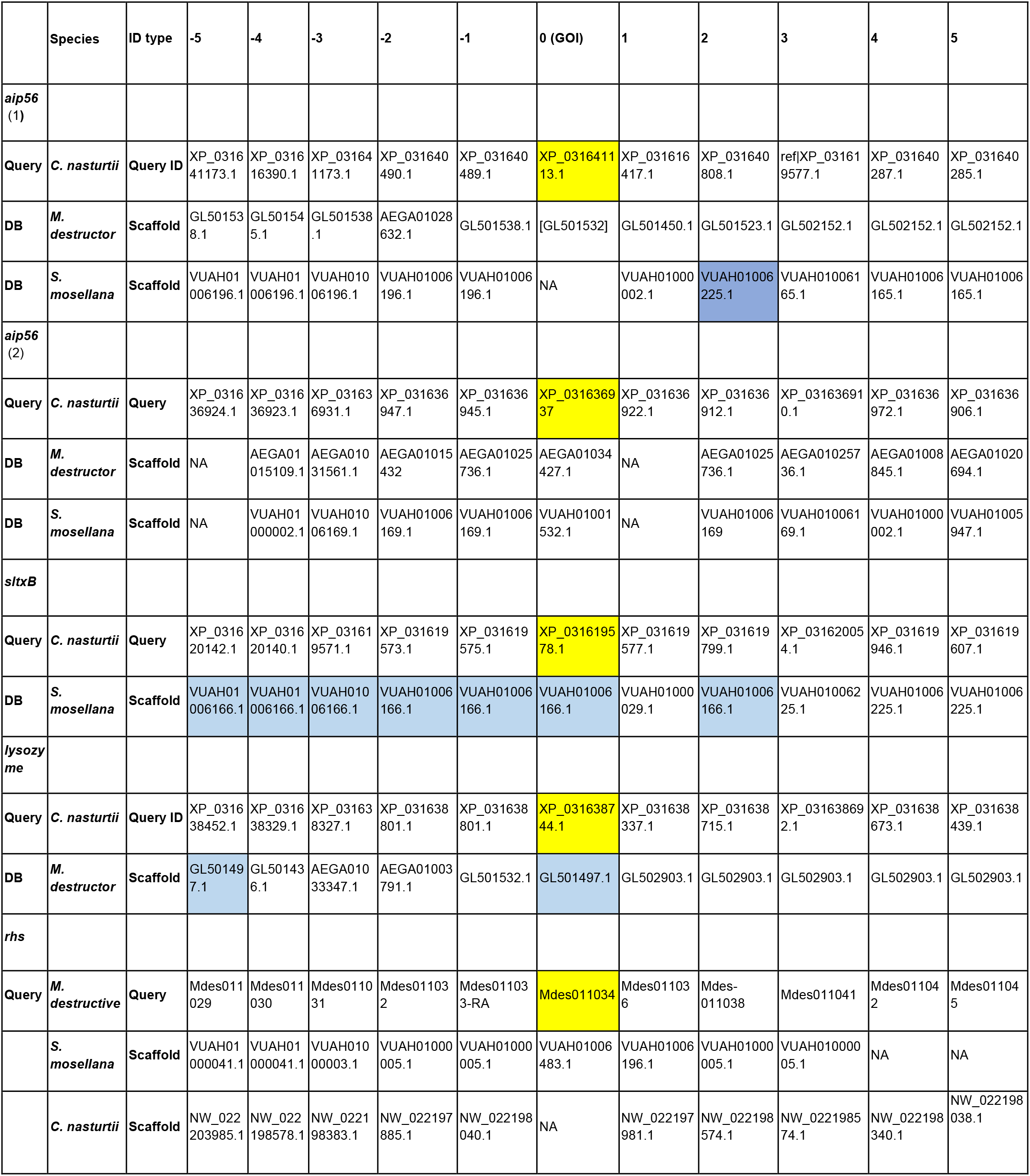
Patterns of micro-synteny show that *sltxB* was likely transferred in Cecidomyiidae prior to the divergence of *C. nasturtii* and *S. mosellana*, but fragmented genomes make the timing of the other horizontally transferred genes difficult to determine. Text highlighted in yellow is the locus (or protein GenBank ID) of interest. “Position relative to GOI” indicates the number of genes upstream or downstream of the gene of interest (GOI). For example, −5 would indicate the gene is five genes upstream of the GOI. Cells highlighted in blue indicate genes are syntenic with the GOI in the considered species.

**Table S4.**
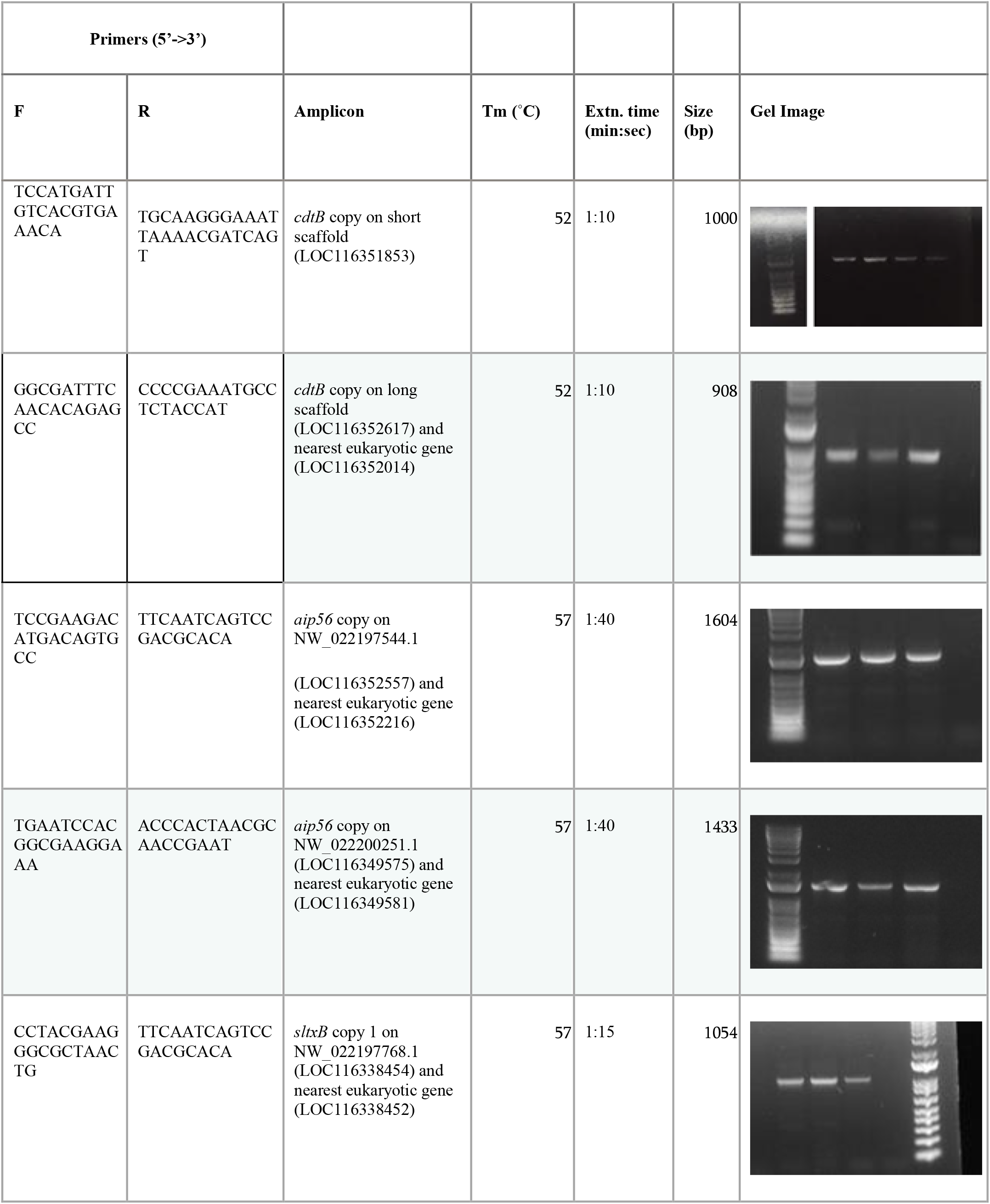
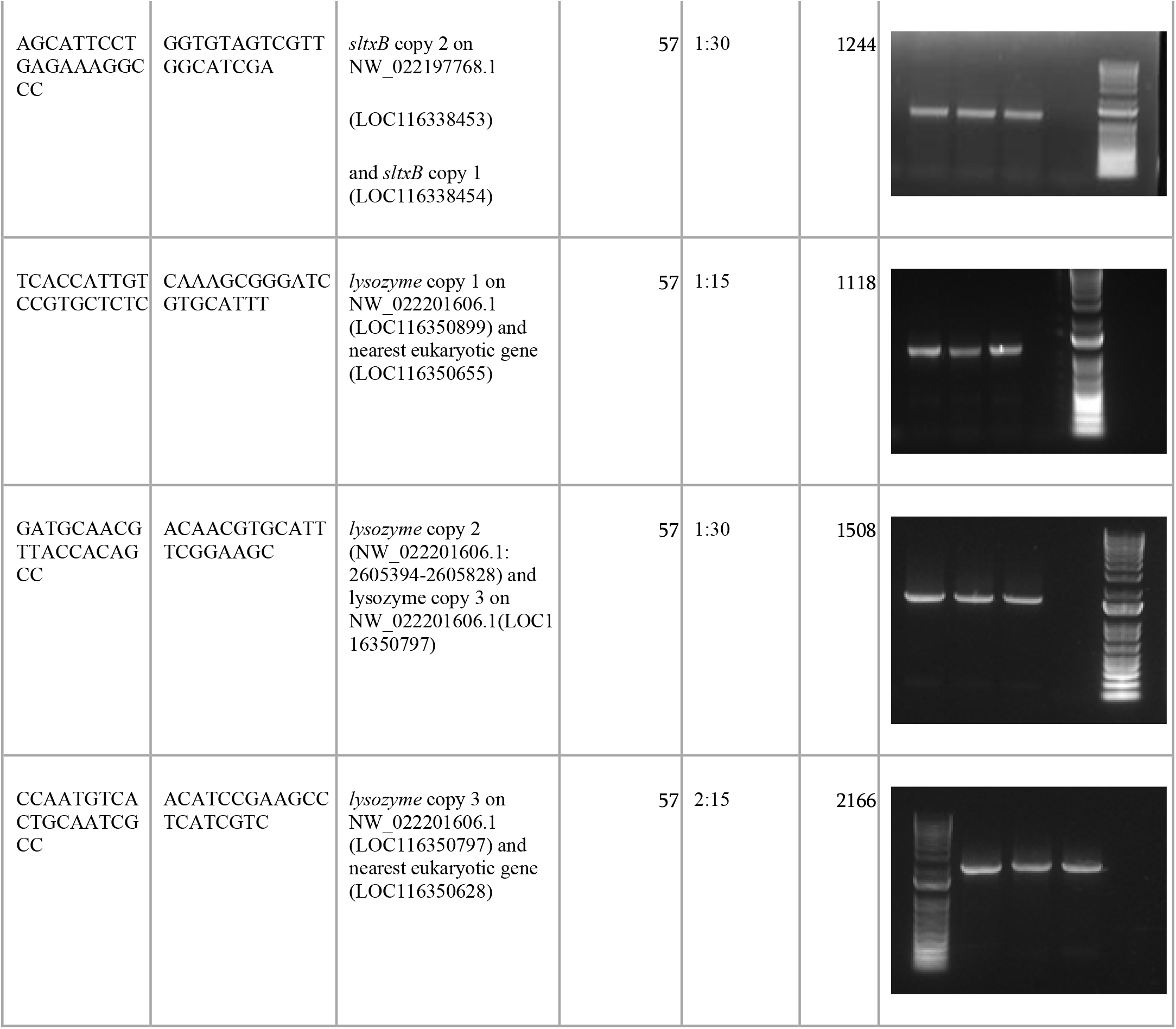
Primers, reaction details, and gel images for amplification of horizontally transferred genes in the *C. nasturtii* nuclear genome.

**Table S5.**
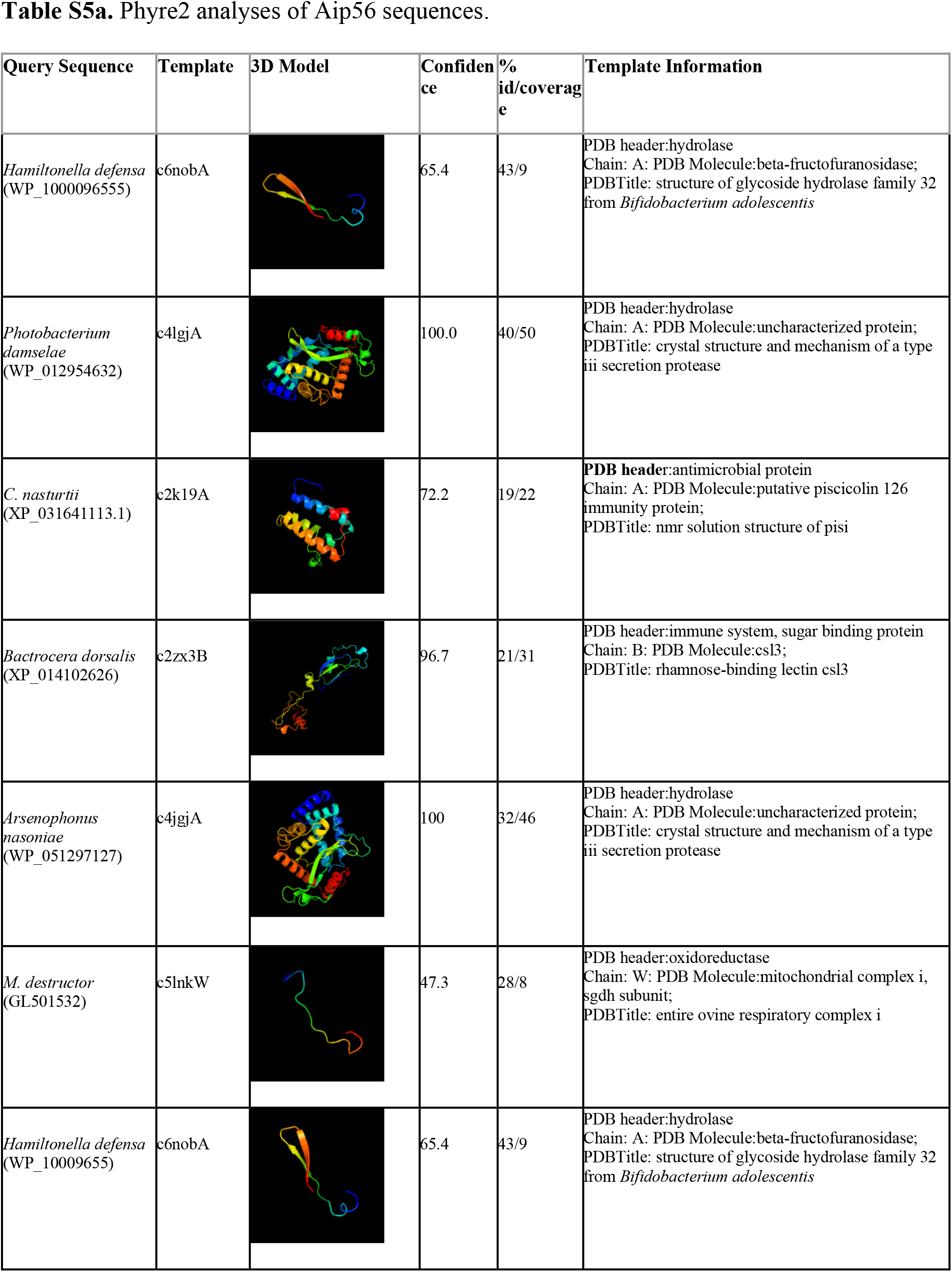
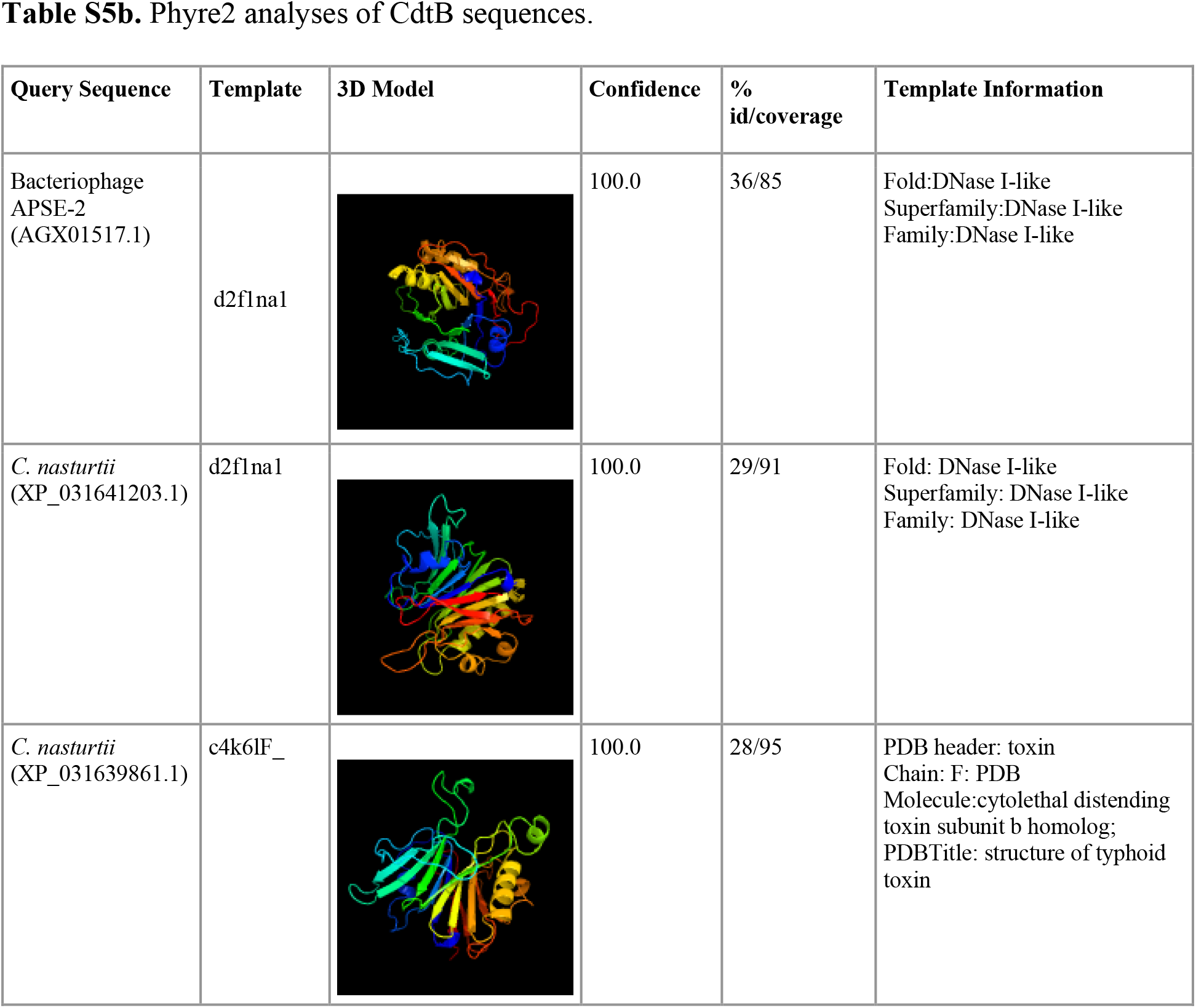
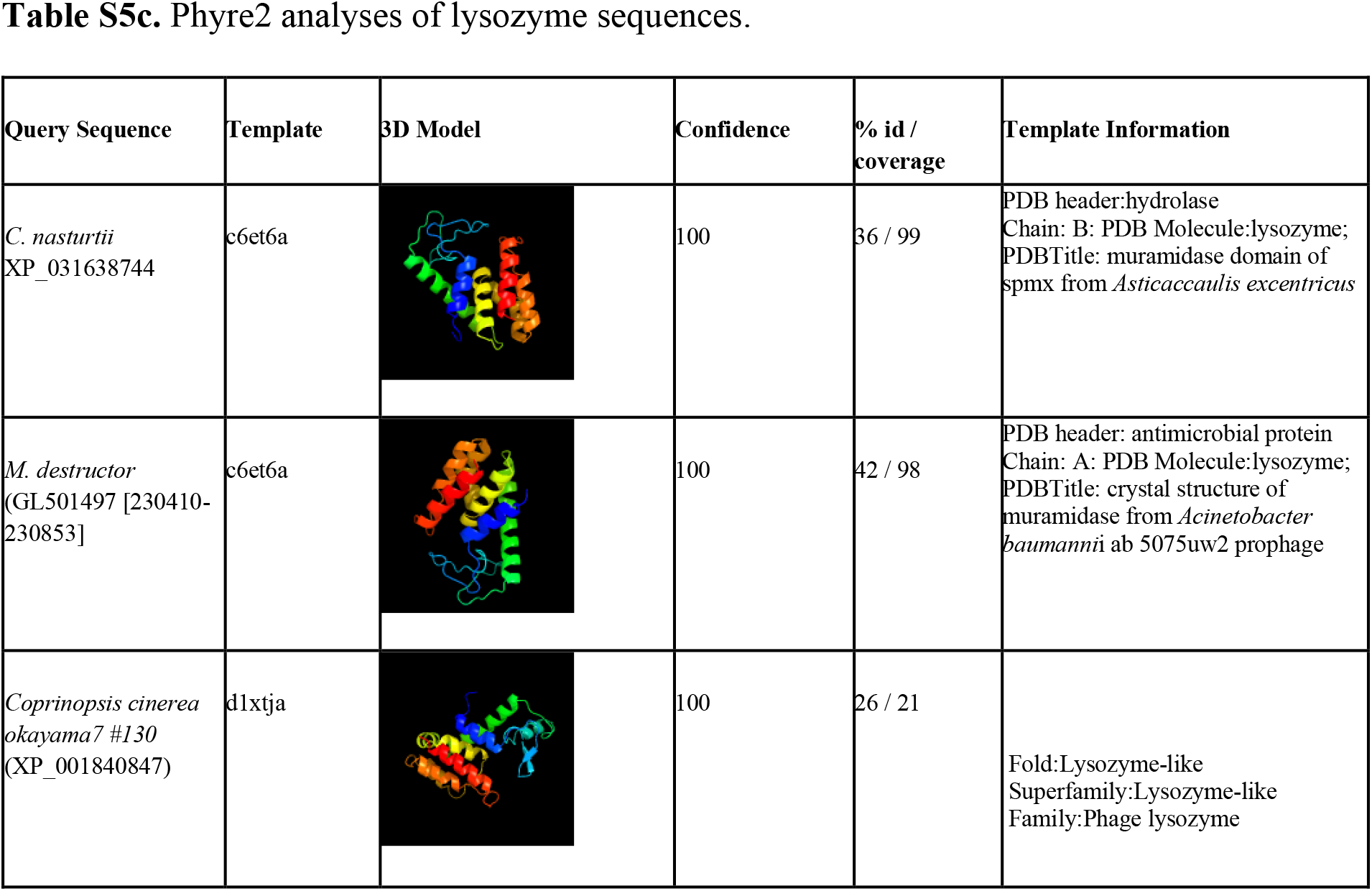
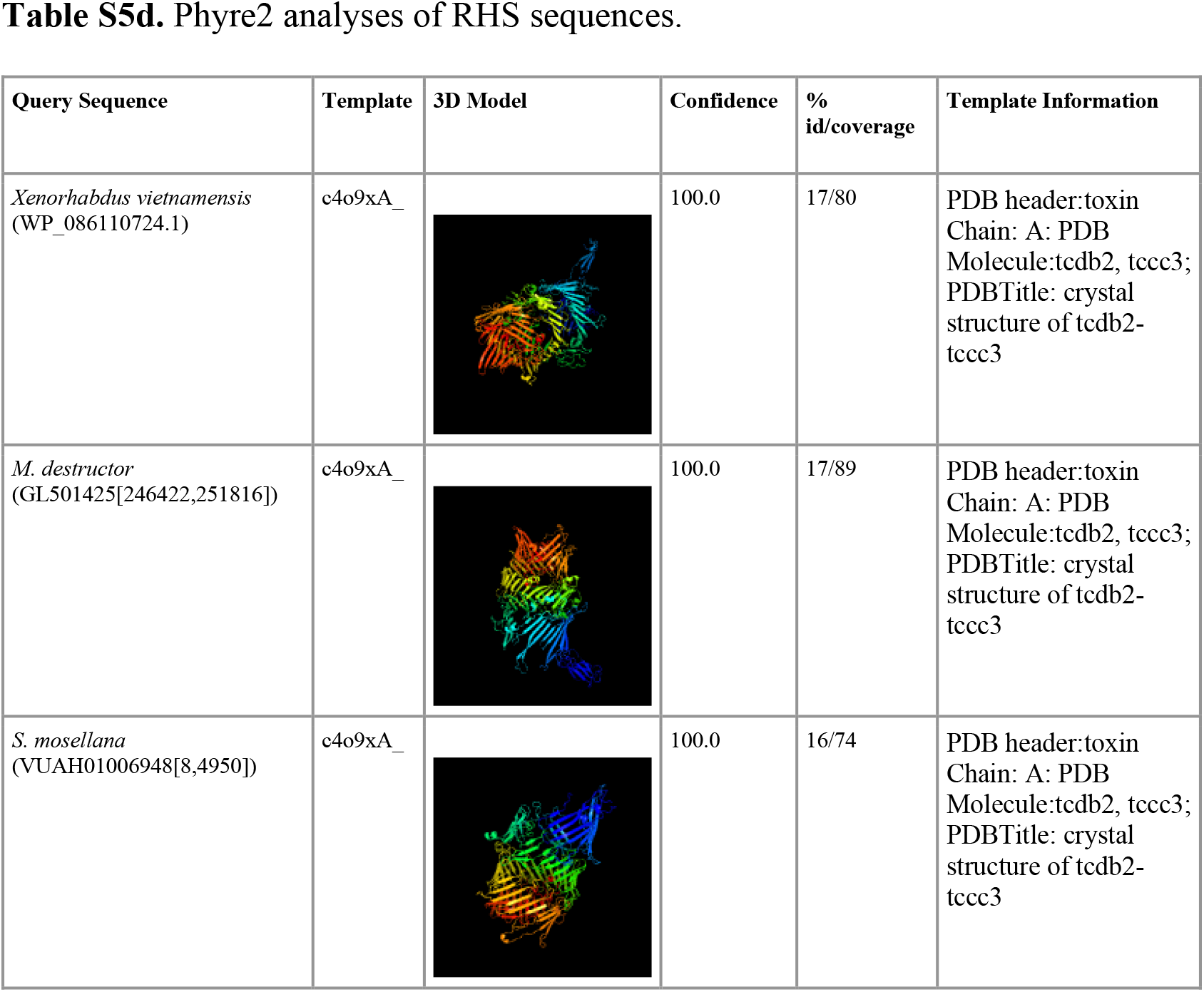
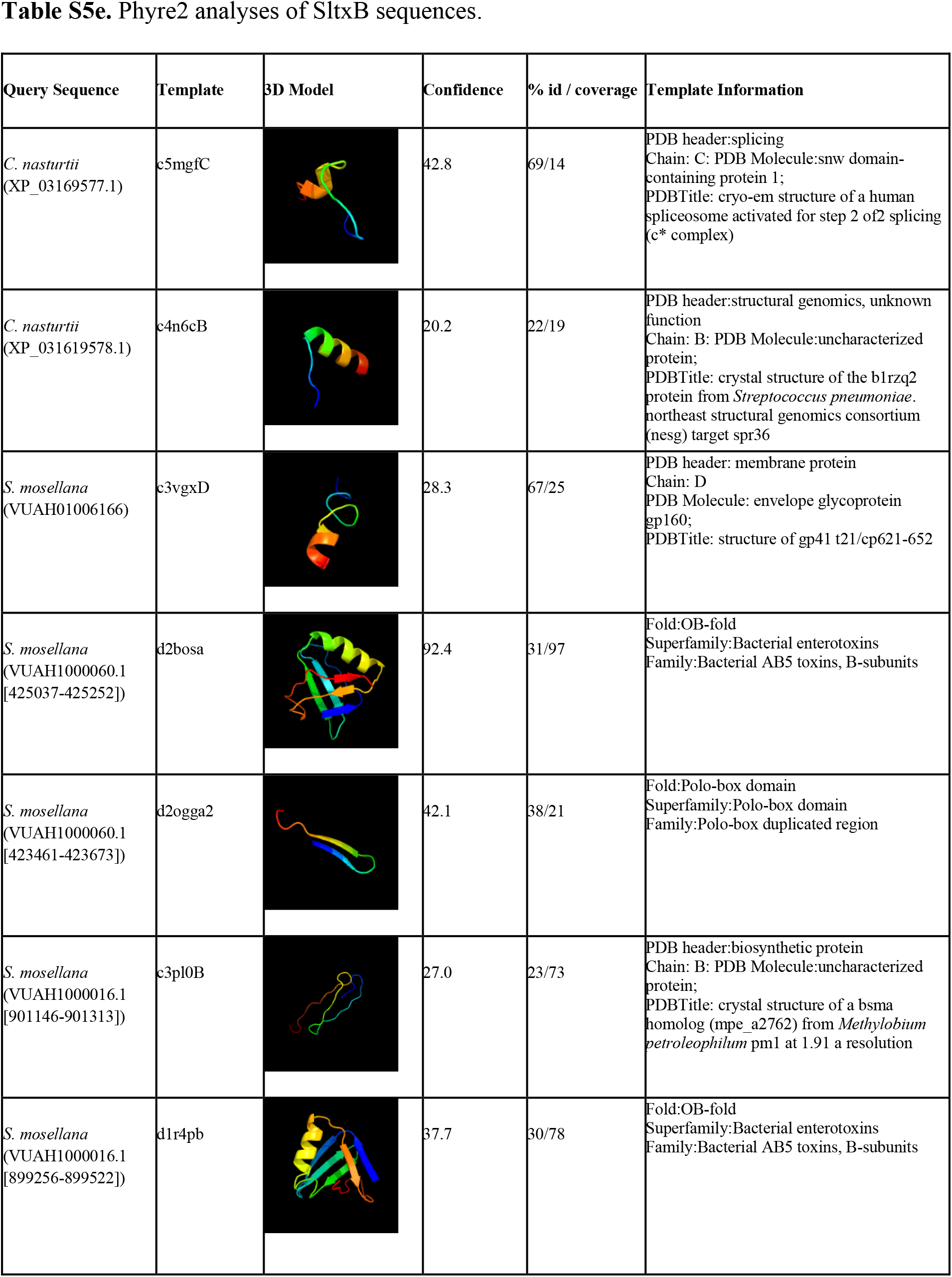
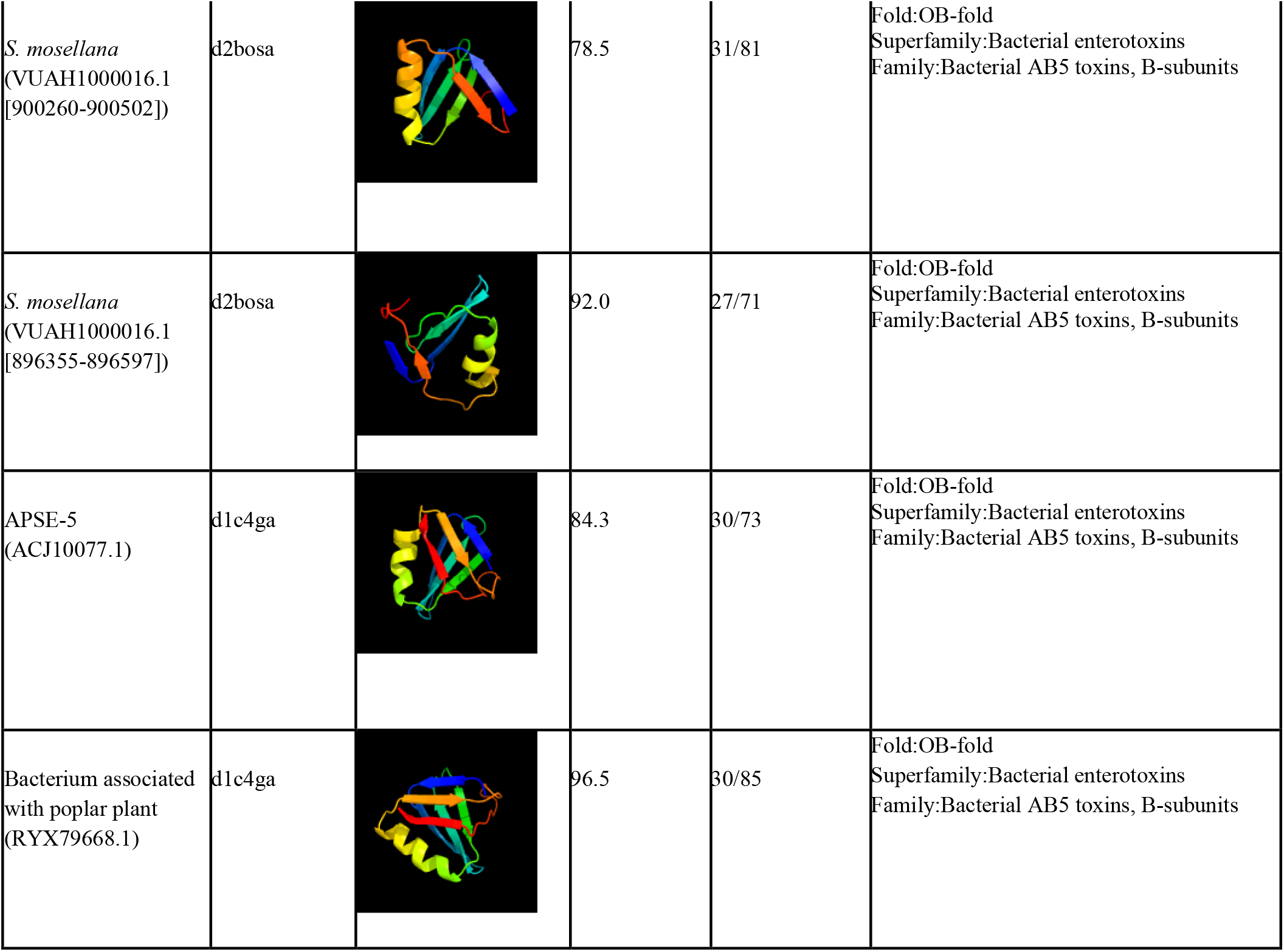
Phyre2 analyses of proteins encoded by horizontally transferred genes can provide clues as to the function of these proteins, even in novel eukaryotic contexts. **Table S5a-e** are Phyre2 analyses for, respectively: Aip56, CdtB, Lysozyme, RHS, and SltxB.

**Table S6.** δ values for gene phylogenies demonstrate that there is a relationship between ecological habitat and horizontal gene transfer. δ values for both complete trees and trees for which we removed vertical descendance (“HGT-Only”) are shown. P-value is calculated as the number of simulations (*n*=100) in which the shuffled δ is higher than the realized δ.

|  | Real Phylogeny |  |  | HGT-Only Phylogeny |  |  |
| --- | --- | --- | --- | --- | --- | --- |
| | Tips | $\delta$ | p-value | Tips | $\delta$ | p-value |
| Aip56 | 90 | 9.55 | <0.01* | 63 | 16.8 | 0.01* |
| SltxB | 28 | 6.86 | <0.01* | 11 | 35.7 | 0.07 |
| CdtB | 76 | 10.3 | <0.01* | 41 | 5.92 | <0.01* |
| RHS | 188 | 4.85 | <0.01* | 109 | 3.98 | <0.01* |
| Lysozyme | 172 | 13.2 | <0.01* | 155 | 13.4 | <0.01* |

**Table S7.** Accession numbers for marker genes included on Cecidomyiidae species tree.

| Species | Genome | Transcriptome | CO1<br>accession # | CAD accession # | EF1a accession # | 28S accession # |
| --- | --- | --- | --- | --- | --- | --- |
| *Bibio marci* * | NA | SRX314826 | KT316846.1 | KX453730.1 | NA | KJ136761.1 |
| *Catotricha subobsoleta* | GCA_011634745.1 | NA | KT316873.1 | KX453747.1 | MG684878.1 | KP288821.1 |
| *C. nasturtii* | GCA_009176525.2 | SRX6853821 | EU812560.1 | XM_031771980.1 | XM_031764621.1 | NA |
| *M. destructor* | GCA_000149185.1 | SRX516926 | EU375697.1 | FJ040625.1 | AF085227.1 | FJ040514.1 |
| *S. mosellana* | GCA_009176505.1 | SRX252524 | KC769209.1 | MN191486.1 | VUAH01006101.1 | MN201531.1 |

\*Outgroup.

**Table S8.**
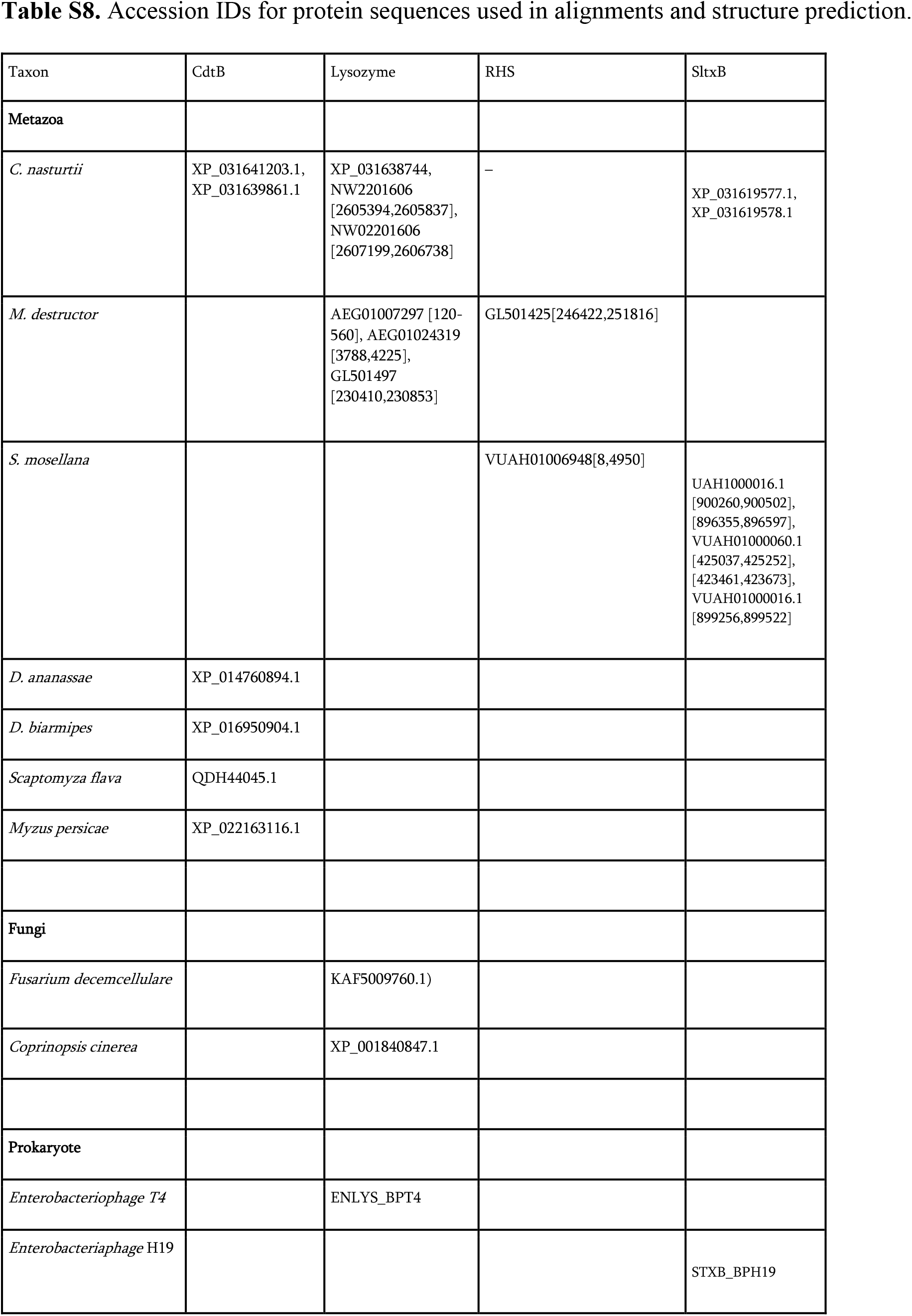
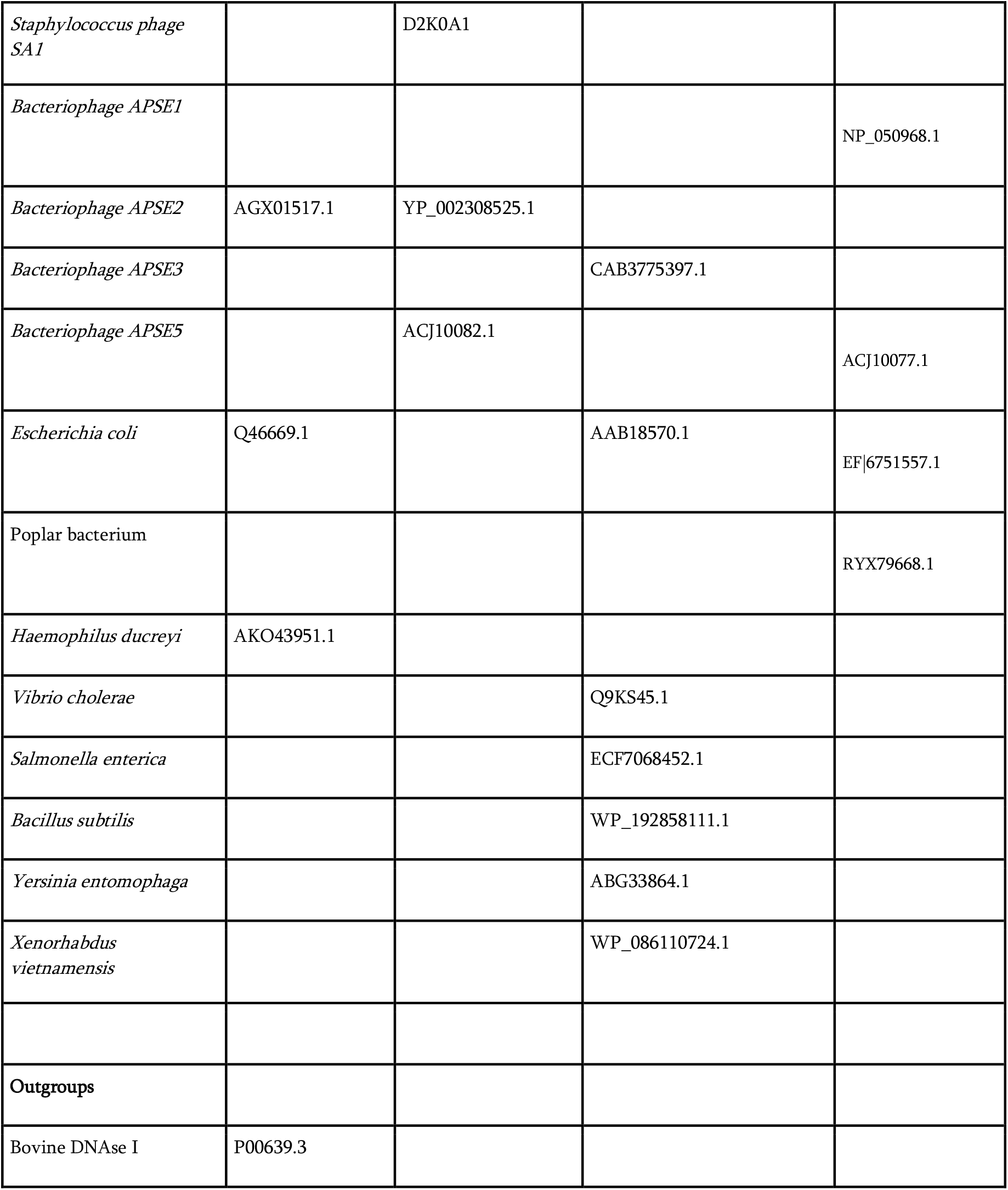
Accession IDs for protein sequences used in alignments and structure prediction.

**Fig S1.**
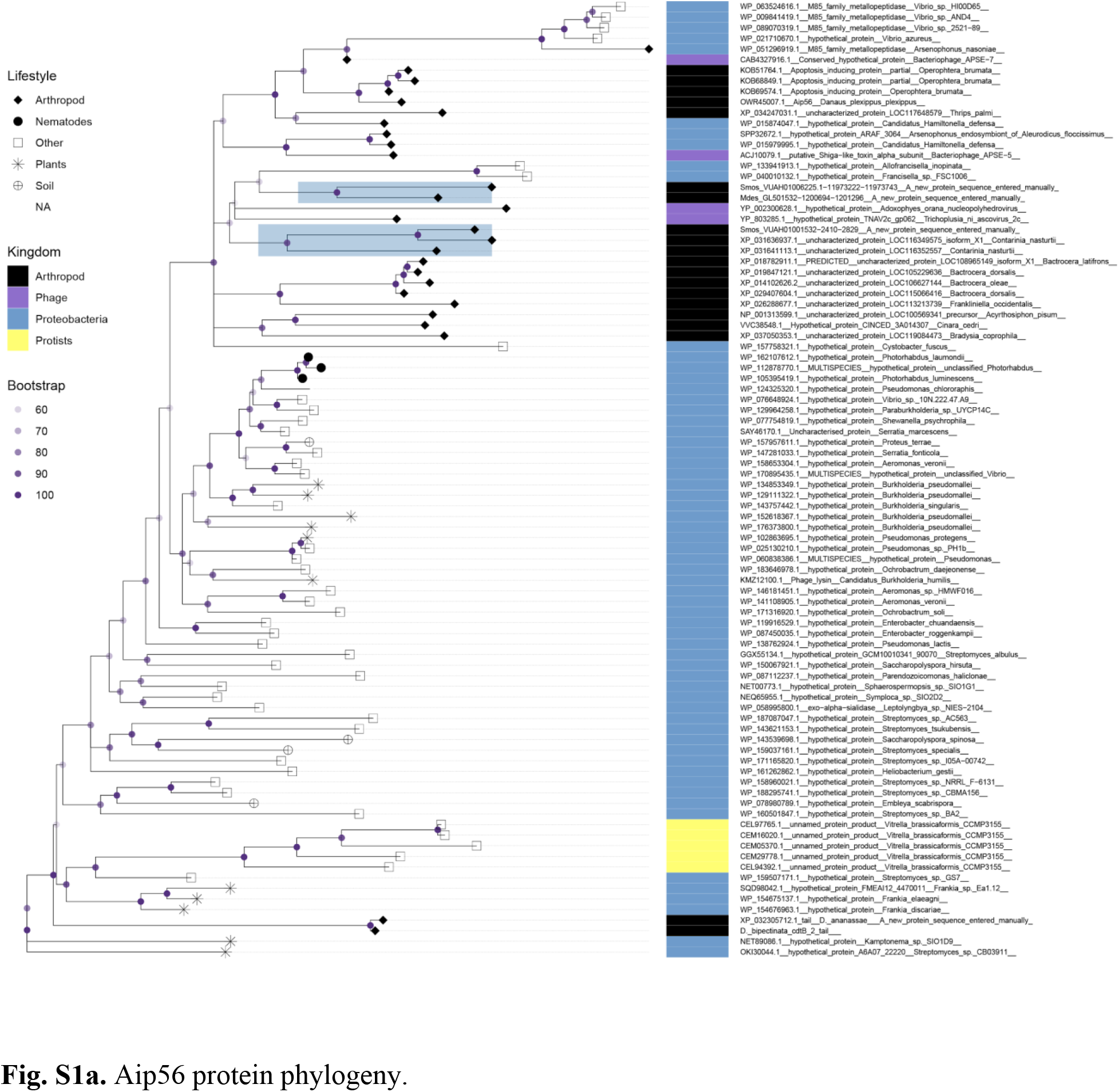
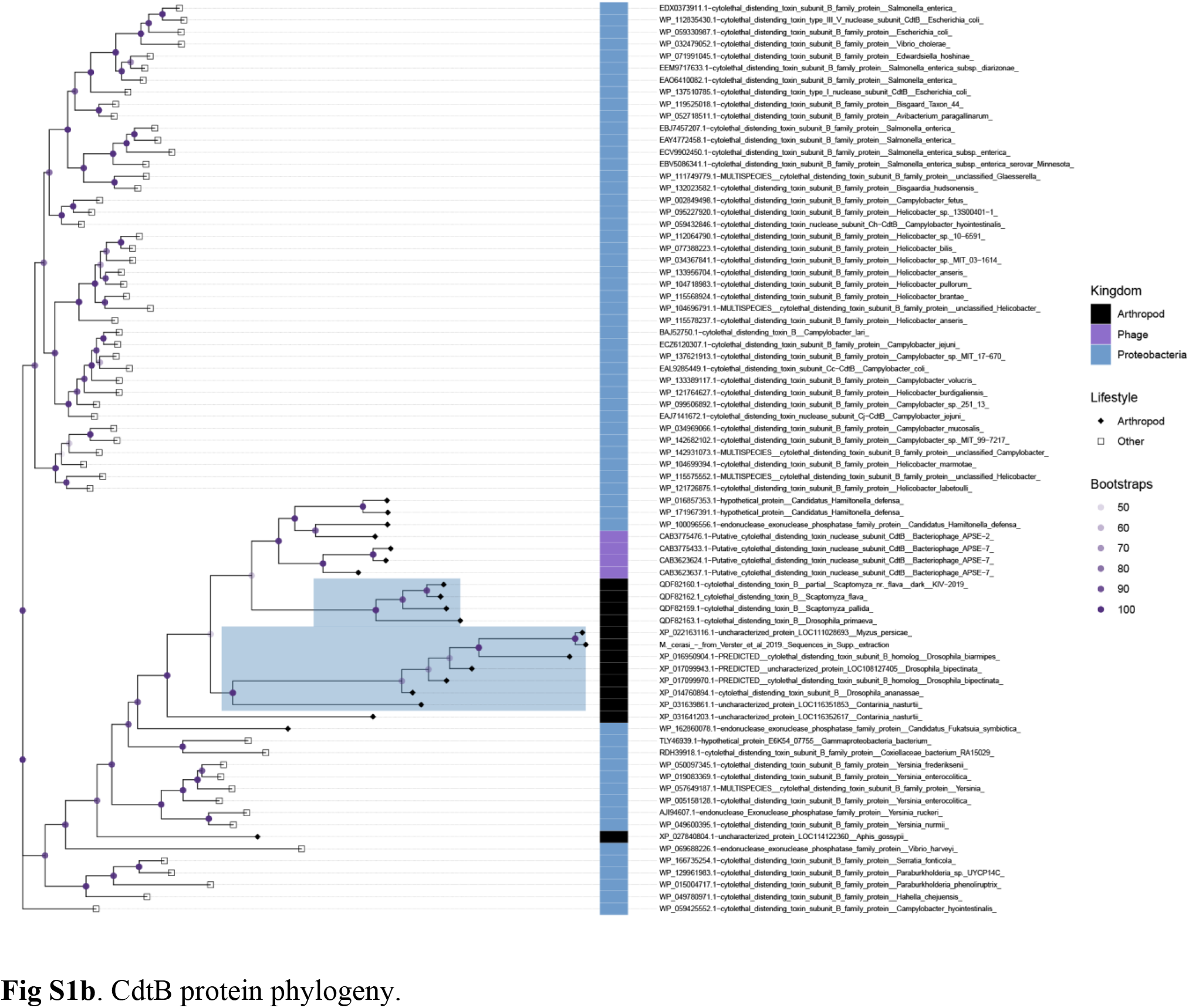
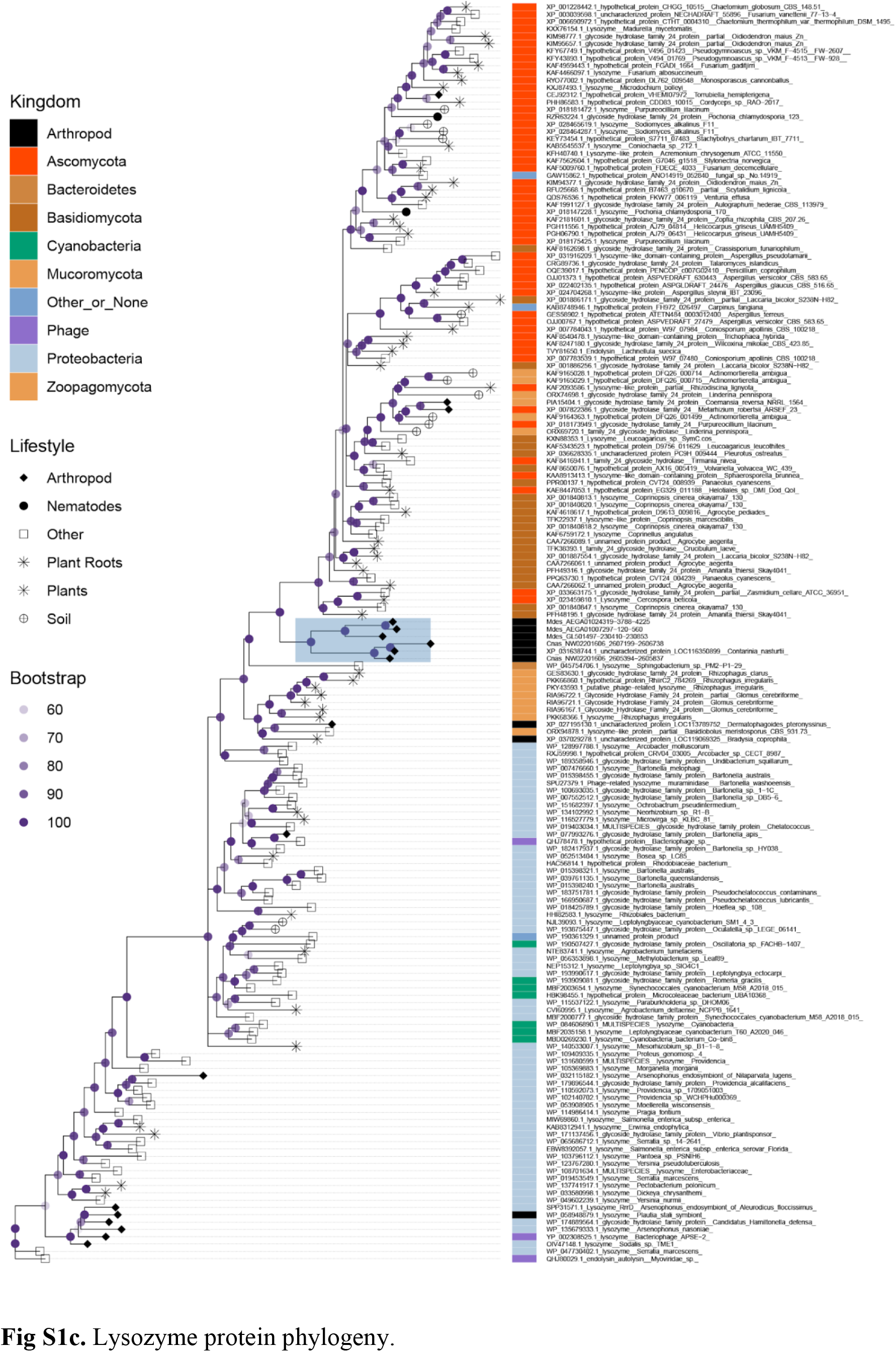
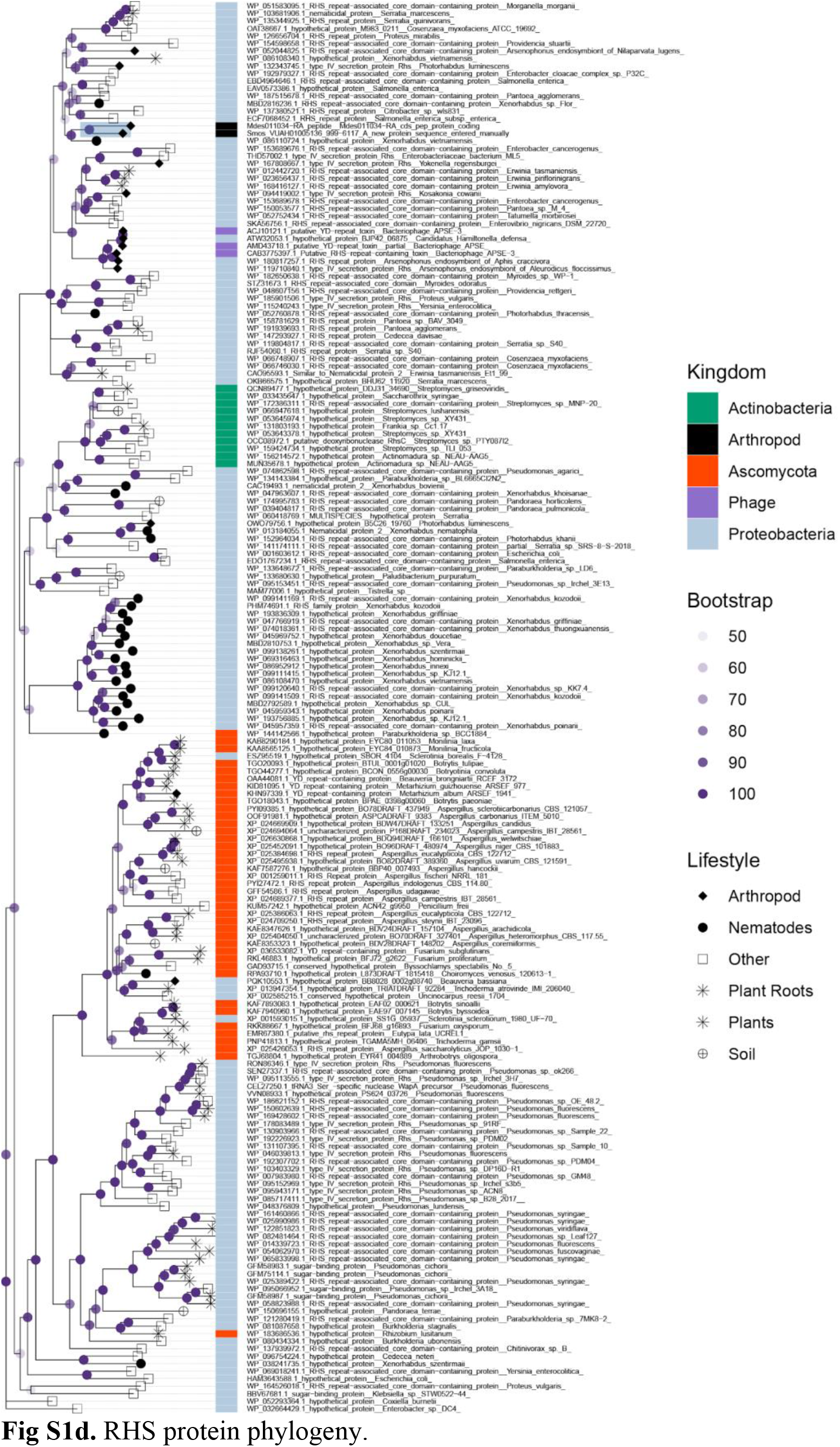
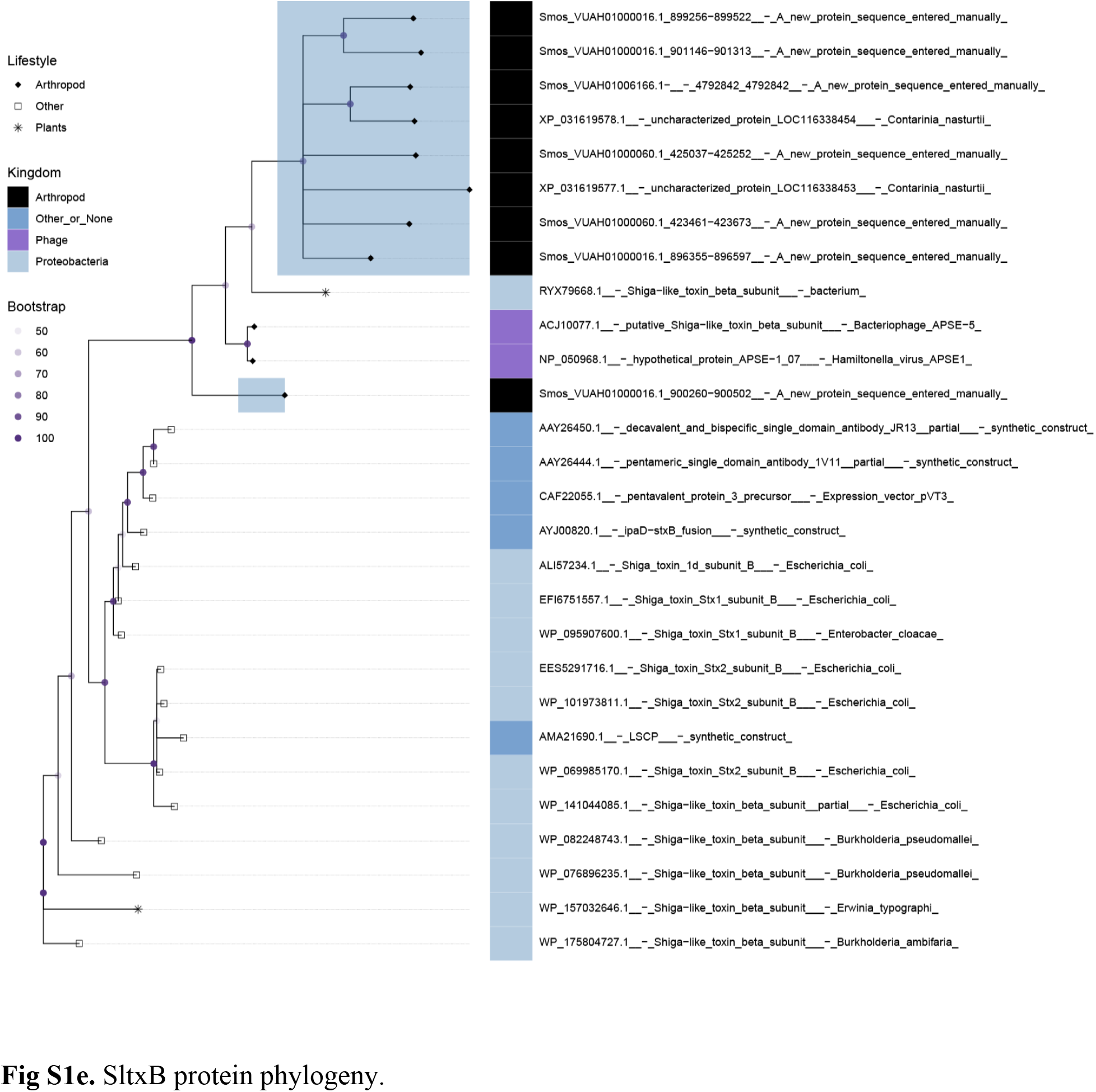
Complete Maximum Likelihood protein phylogenies for five genes transferred to the Cecidomyiidae lineage reveal possible donor provenance. Phylogenies show taxonomic and lifestyle information for each tip. HTG clades discussed in the manuscript are highlighted in blue. Branch bootstrap values are shown as darkness of nodes and are based on 1000 ultrafast bootstrap replicates. **Figs S1a-e** show, respectively: Aip56, CdtB, Lysozyme, RHS, and SltxB. For more information about the phylogenies, see **Supplementary Methods**.

**Fig S2.**
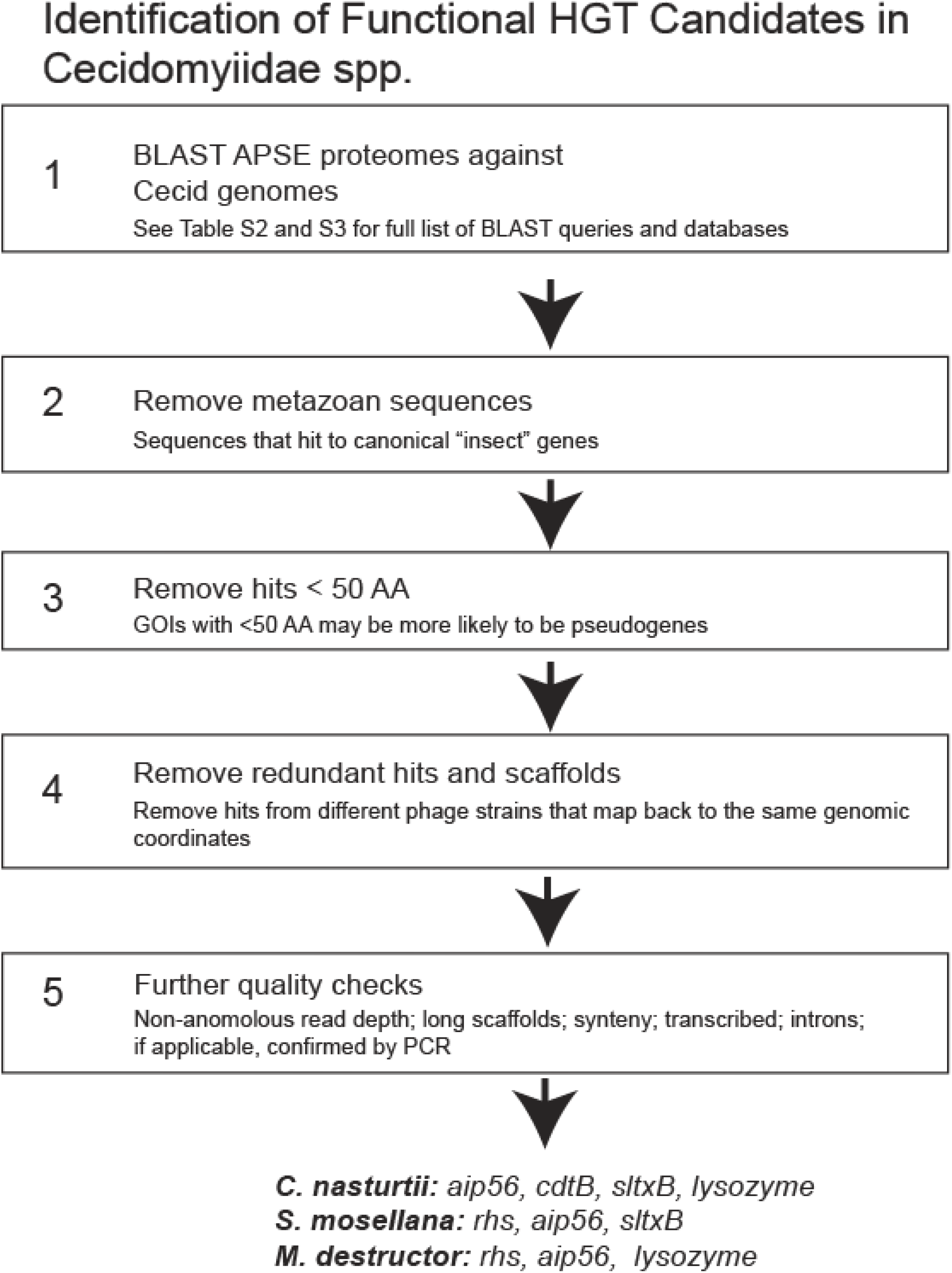
Workflow to identify functional horizontal gene transfer candidates in Cecidomyiidae species identifies at least five transfers of toxin-encoding genes.

**Fig S3.**
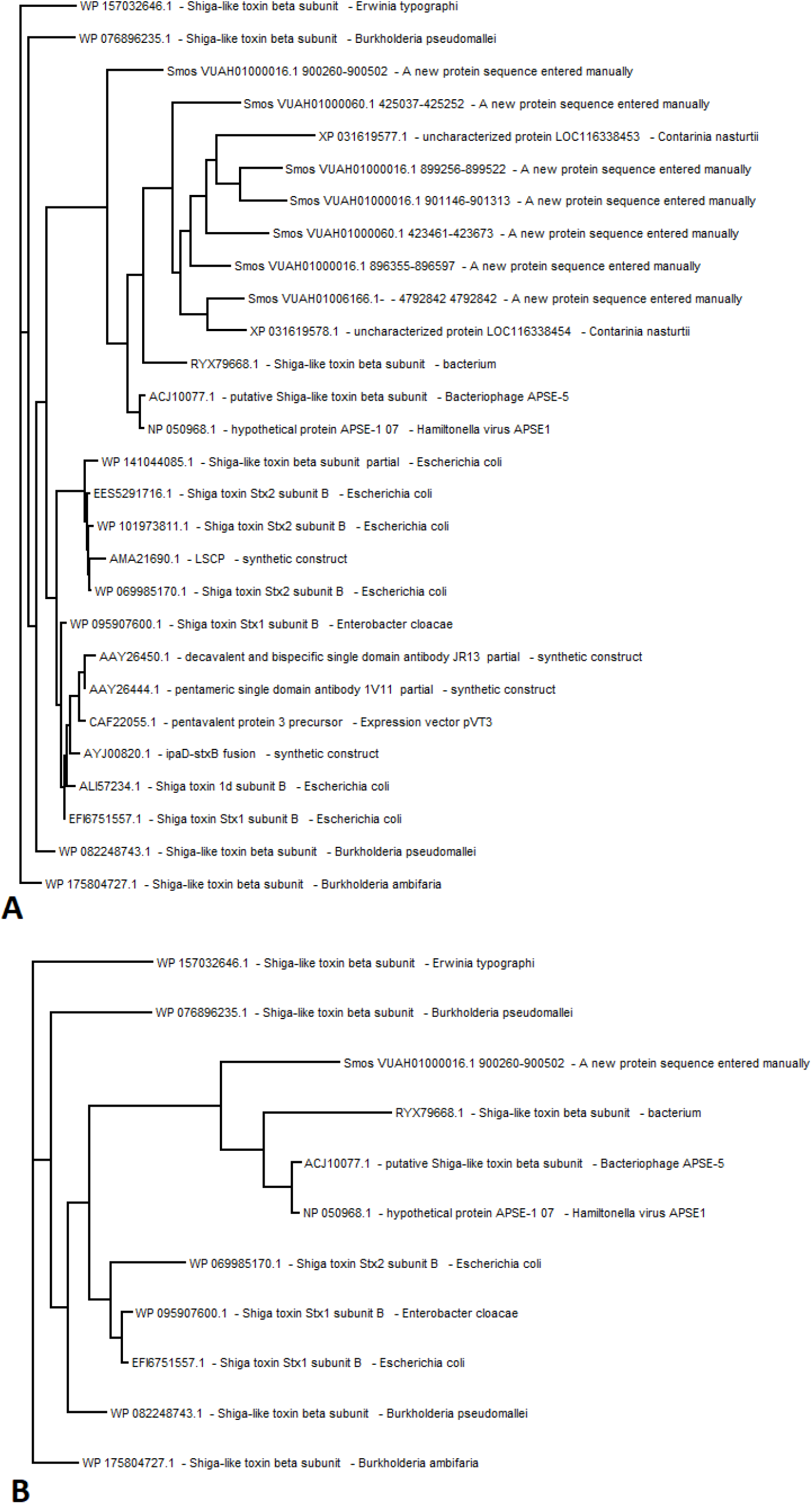
Two representative trees that were used in the comparison of phylogenetic signal. The tree labeled ‘A’ is the un-pruned tree, while the tree shown in ‘B’ has been trimmed to only include tips that were likely passed down via horizontal transfer.

**Supplementary File 1 Legend**. These analyses support the finding that the horizontally transferred genes identified in this study are not due to contamination. ‘Species’ column shows the species in which the HGT occurred. ‘APSE’ and ‘Protein ID’ columns show the APSE strain and GenBank IDs of query sequences identified in the cecidomyiid genomes. For the ‘Protein Name’ column, we report a summary of the BLASTP results if they appear to correspond to one or more characterized proteins. In some cases, the identity and function of the protein are ambiguous and are labelled as ‘Hypothetical Protein.’ In the ‘E-value’ column, we report the lowest E-value in the case of multiple APSE protein queries. In some cases, a single TBLASTN query resulted in hits to multiple genomic ‘ranges’ on the same scaffold. If the subject sequences shared high AA identity **(>90%)** throughout multiple ranges, we considered these evidence of duplications of the GOI, and the E-values for each individual ‘range’ was reported in separate rows. ‘Scaffold’ and ‘Scaffold Size’ coordinates reflect GenBank accessions and associated lengths unless otherwise noted. ‘GOI Coordinates’ column reflects the TBLASTN reported ranges, unless the GOI has been annotated, in which case the annotation ID is shown. In the ‘Other Eukaryotic Genes’ column, we report if we found evidence of *bona fide* eukaryotic genes (Yes/No). Where the genome has been annotated, we report the Annotation IDs of the nearest eukaryotic gene proximal to the GOI. If the genome has not been annotated, we ran Augustus annotation on each scaffold under consideration using the ‘fly’ setting as implemented in Geneious (Stanke et al. 2004). In the ‘Intron’ and ‘Exon Coordinates’ column, we indicate the number of introns predicted by either annotations specific to the species or Augustus annotations. In some cases, Augustus did not predict any genes in the region of interest, in which case we reported ‘NGP’ for ‘No Gene Predicted.’ Note that Augustus relies on training on the appropriate gene sets (Stanke et al. 2004), and it may fail in cases of HGT due to the inherent differences of genes with lateral provenance. Where the GOI does not have an associated annotation ID, we report the Augustus-predicted exon coordinates. For the ‘BWA’ columns, please see **Supplementary Methods** - *BWA analysis* section. For ‘Transcription’ columns, please see **Supplementary Methods** - *Transcription analysis*. In the case of *C. nasturtii*, we indicate if we were able to successfully PCR the GOI (PCR primers and conditions are documented in **Table S4**).

**Supplementary File 2 Legend**. Taxonomic and lifestyle information for species in protein phylogenies shown in **Fig S1**. Included are citations if species are found on Arthropods, Plants or Soil.

